# Syntrophy via interspecies H_2_ transfer between *Christensenella* and *Methanobrevibacter* underlies their global co-occurrence in the human gut

**DOI:** 10.1101/872333

**Authors:** Albane Ruaud, Sofia Esquivel-Elizondo, Jacobo de la Cuesta-Zuluaga, Jillian L. Waters, Largus T. Angenent, Nicholas D. Youngblut, Ruth E. Ley

## Abstract

Across human populations, 16S rRNA gene-based surveys of gut microbiomes have revealed that the bacterial family *Christensenellacea*e and the archaeal family *Methanobacteriaceae* co-occur and are enriched in individuals with a lean, compared to an obese, BMI. Whether these association patterns reflect interactions between metabolic partners remains to be ascertained, as well as whether these associations play a role in the lean host phenotype with which they associate. Here, we validated previously reported co-occurrence patterns of the two families, and their association with a lean BMI, with a meta-analysis of 1,821 metagenomes derived from 10 independent studies. Furthermore, we report positive associations at the genus and species level between *Christensenella* spp. and *Methanobrevibacter smithii,* the most abundant methanogen of the human gut. By co-culturing three *Christensenella* spp. With *M. smithii,* we show that *Christensenella* spp. efficiently support the of *M. smithii* via H_2_ production, far better than *Bacteroides thetaiotaomicron*. *C. minuta* forms flocs colonized by *M. smithii* even when H_2_ is in excess. In culture with *C. minuta*, H_2_ consumption by *M. smithii* shifts the metabolic output of *C. minuta*’s fermentation towards acetate rather than butyrate. Together, these results indicate that the widespread co-occurrence of these microbiota is underpinned by both physical and metabolic interactions. Their combined metabolic activity may provide insights into their association with a lean host BMI.

**Importance:** The human gut microbiome is made of trillions of microbial cells, most of which are *Bacteria*, with a subset of *Archaea*. The bacterial family *Christensenellaceae* and the archaeal family *Methanobacteriaceae* are widespread in human guts. They correlate with each other and with a lean body type. Whether species of these two families interact, and how they affect the body type, are unanswered questions. Here, we showed that species within these families correlate with each other across people. We also demonstrated that particular species of these two families grow together in dense flocs, wherein the bacteria provide hydrogen gas to the archaea, which then make methane. When the archaea are present, the ratio of bacterial products (which are nutrients for humans) is changed. These observations indicate when these species grow together, their products have the potential to affect the physiology of their human host.

## Introduction

Obesity was the first human disease phenotype to be associated with an altered microbial ecology of the gut (1, 2). The link between the relative abundance in the gut of the bacterial family *Christensenellaceae* and a low host body mass index (BMI) now stands as one of the most robust associations described between the human gut microbiome and host BMI (3–15). Compared to other families of bacteria that comprise the human gut microbiome, the family *Christensenellaceae* was described relatively recently, when the type strain *Christensenella minuta* was reported in 2012 (16). Prior to the description of *C. minuta*, 16S rRNA sequences from this genus escaped notice in the gut microbiome, though these sequences accumulated steadily in SSU rRNA gene databases. A positive association between a lean host BMI and the relative abundance in the gut of *Christensenellaceae* 16S rRNA genes was first reported in 2014 (4). The association was shown to have existed in earlier datasets (4), but was likely undetected as this family had not yet been named. Goodrich *et al.* showed a causal link between the *Christensenellaceae* and host BMI in gnototioc mice: the addition of *C. minuta* to the gut microbiome of an obese human donor prior to transplantation reduced adiposity gains in the recipient mice compared to controls receiving the unsupplemented microbiome (4). The mechanism underlying this host response remains to be elucidated. One step towards this goal is a better understanding of how the members of the *Christensenellaceae* interact ecologically with other members of the gut microbiome.

Across human populations, gut microbiota often form patterns of co-occurrence (*e.g.*, when these consortia exist in a subset of human subjects, they are termed enterotypes (17)). Such co-occurrences of taxa across subjects reflect shared environmental preferences, but to determine if they represent metabolic or physical interactions requires further study. The family *Christensenellaceae* consistently forms the hub of co-occurrence networks with other taxa (6, 8, 9, 18, 19). Notably, gut methanogens (specifically, of the archaeal family *Methanobacteriaceae*) are often reported as part of the *Christensenellaceae* co-occurrence consortium (4, 20–22). The most widespread and abundant of the gut methanogens, *Methanobrevibacter smithii*, produces CH_4_ from H_2_ and CO_2_, the products of bacterial fermentation of dietary fibers. Such cross-feeding likely explains why the relative abundances of *M. smithii* and fermenting bacteria are often positively correlated (21, 23, 24). Several studies have shown that in the laboratory, *M. smithii* can grow from the H_2_ provided by *Bacteroides thetaiotaomicron*, a common gut commensal bacterium (25–27). Given that the cultured representatives of the *Christensenellaceae* ferment simple sugars (16, 28), and that their genomes contains hydrogenases (29), we predicted that members of the *Christensenellaceae* produce H_2_ used by *M. smithii* as a substrate in methanogenesis.

Here, we explored the association between the *Christensenellaceae* and the *Methanobacteriaceae* in two ways. First, we analyzed metagenomes for statistical associations between the two families and their subtaxa. Compared to 16S rRNA gene surveys, metagenomes often can better resolve the taxonomic assignments of sequence reads below the genus level (30). Metagenome-based studies have so far been blind to the *Christensenellaceae*, however, because their genomes have been lacking from reference databases. Here, we customized a reference database to include *Christensenellaceae* genomes, which we used in a meta-analysis of >1,800 metagenomes from 10 studies. Second, to assess for metabolic interactions between members of the *Christensenellaceae* and *Methanobacteriaceae,* we measured methane production by *M. smithii* when grown in co-culture with *Christensenella* spp. Our results show that: i) the positive association between the *Christensenellaceae* and the *Methanobacteriaceae* is robust to the genus/species level across multiple studies; ii) these taxa associate with a lean host BMI; iii) *Christensenella* spp. support the growth of *M. smithii* by interspecies H_2_ transfer far better than *B. thetaiotaomicron*; and iv) *M. smithii* directs the metabolic output of *C. minuta* towards less butyrate and more acetate and H_2_, which is consistent with reduced energy availability to the host and consistent with the association with a low BMI.

## Results

### *Christensenella* relative abundance is significantly correlated with leanness across populations

Both the *Christensenellaceae* family and the genus *Christensenella* had a very high prevalence, as they were present in more than 99% of the 1,821 samples; both the family and the genus have a mean abundance of 0.07% ± 0.05 (Fig. 1b,d and Fig. S1). To correct for the influence of environmental factors on the relative abundance of the *Christensenellaceae* family and of the *Christensenella* genus, we first constructed null models in which we selected covariates (Appendix 1) that explained a significant proportion of the variance of the transformed relative abundance of the family *Christensenellaceae*, and in the same manner of the *Christensenella* genus. BMI and age were significantly correlated to the transformed relative abundances of *Christensenellaceae* and of *Christensenella* (designated with the suffix ‘-tra’, *i.e.*, Cf-tra and Cg-tra), and were retained in the null models (Cf-null and Cg-null).

**Fig. 1.**
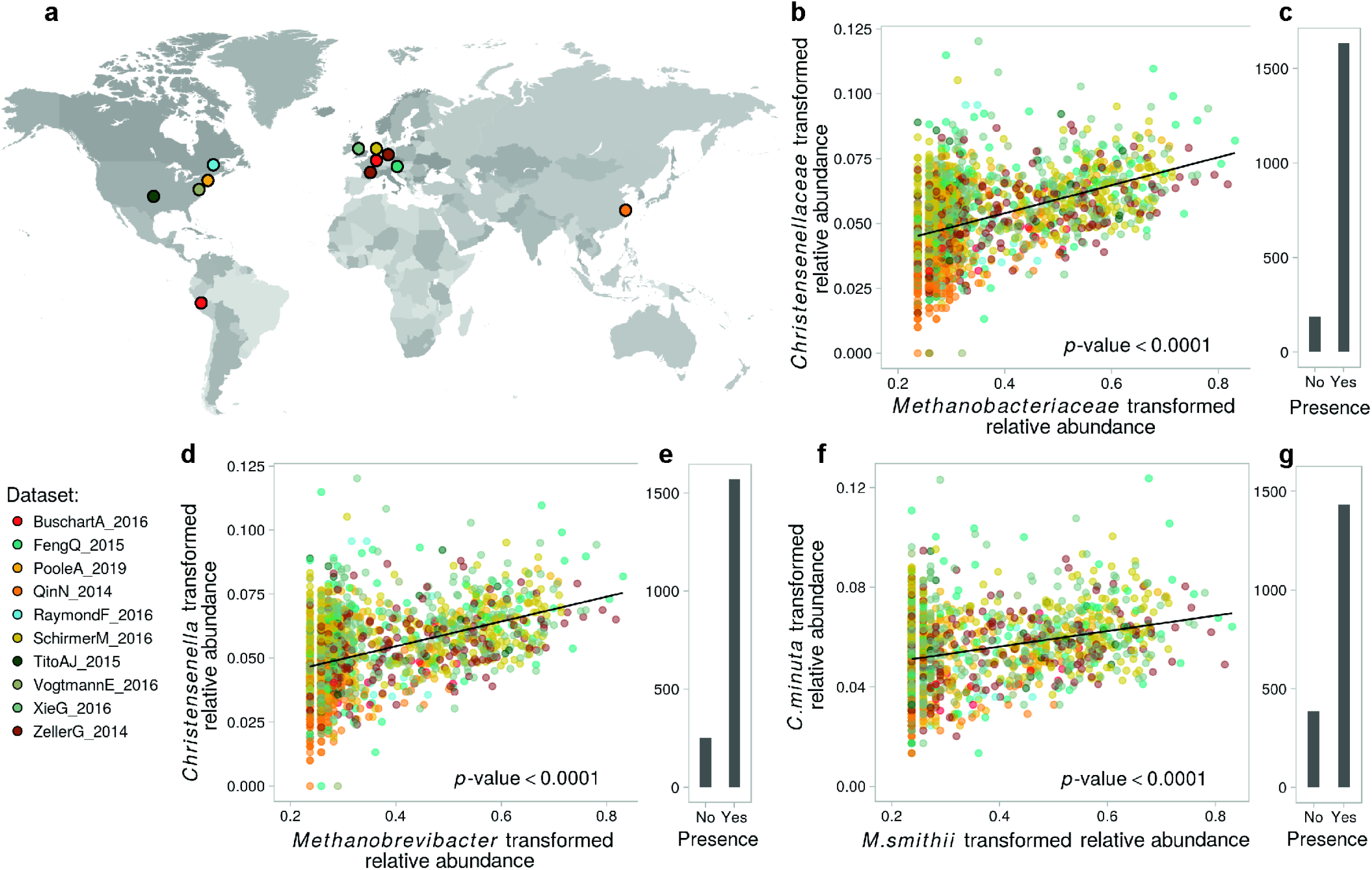
Abundances of the *Methanobacteriaceae* and *Christensenellaceae* families across populations. (a) Countries where the human gut metagenomes used in our meta-analysis (n = 1,821 samples) were recruited by 10 independent studies (summarized in Dataset); (b) association between the transformed relative abundances of *Christensenellaceae* and *Methanobacteriaceae*, in samples where the *Methanobacteriaceae* was detected; (c) Number of samples for which the *Methanobacteriaceae* were detected; d-e and f-g: same as b-c, at the genus and species level respectively. The correlation between the transformed relative abundances of both taxa at each taxonomic level was evaluated using linear mixed models to corrected for covariates (ANOVA, p-values < 0.0001).

BMI was negatively correlated to both Cf-tra (type II ANOVA, p-value = 0.0002 and F-value(339) = 14.46) and Cg-tra (type II ANOVA, p-value = 0.0002 and F-value(339) = 14.29), indicating that leaner individuals harbor higher relative abundances of *Christensenellaceae* and *Christensenella*. Age was negatively correlated with Cf-tra (type II ANOVA, p-value = 0.01 and F-value(1, 468) = 6.56) and with Cg-tra (type II ANOVA, p-value = 0.01 and F-value(1, 468) = 6.53), indicating that younger subjects carry a greater relative abundance of *Christensellaceae* and *Christensenella*. However, the interaction term between BMI and age was not significantly correlated to the transformed relative abundances (type II ANOVA, p-values > 0.1), indicating that their effects are additive. These results show that regardless of their BMI, younger subjects have higher levels of *Christensenellaceae* and *Christensenella*, and that the lower a subject’s BMI, the more of these microbes they harbor, regardless of their age.

### *Methanobrevibacter* relative abundance is significantly correlated with leanness across populations

The *Methanobacteriaceae* family and *Methanobrevibacter* genus also had a high prevalence with 92% and 89% of the people harboring them respectively, with mean abundances of 0.48% ± 1.55 and 0.49% ± 1.54, respectively. As above, we evaluated the association between the *Methanobacteriaceae* family, and of the *Methanobrevibacter* genus, with BMI and age (models Mf-null and Mg-null). The transformed relative abundances of *Methanobacteriaceae*, Mf-tra, and of *Methanobrevibacter*, Mg-tra, were also negatively correlated to BMI (type II ANOVA, respective p-values = 0.01 and 0.02, F-values(341; 341) = 6.66 and 5.11). In contrast to the *Christensenellaceae*, both *Methanobacteriaceae* and *Methanobrevibacter* were positively correlated with age (type II ANOVA, respective p-values = 0.001 and 4.27×10^-4^, F-values(1,468; 1,468) = 10.35 and 12.47), indicating that older people carry a greater proportion of methanogens. Moreover, *M. smithii*, the most abundant and prevalent methanogen species within the human gut, was also positively correlated with age and negatively with BMI regardless of age, *i.e.* the interaction term between age and BMI was not significantly correlated (Appendix 2, additional statistics).

### The relative abundances of the *Christensenella* and *Methanobrevibacter* genera are significantly correlated across populations

Next, we looked into how the *Christensenellaceae* and the *Methanobacteriaceae* correlated with each other across subjects while controlling for BMI and age. We constructed a model where Mf-tra was included in addition to BMI and age (model Cf-Mf). This allowed us to test whether adding Mf-tra to the model improved its fit and if so, how much of the variance of Cf-tra not explained by age and BMI could be explained by Mf-tra. We also evaluated the interaction terms between Mf-tra and BMI, and between Mf-tra and age, to assess whether the correlation between Cf-tra and Mf-tra was dependent on age and BMI. The interaction term for BMI and Mf-tra was not significant and was removed from the model; the interaction term for age and Mf-tra was significant and was retained (type I ANOVA, F-value(339) = 8.30 and p-value = 0.0042). We compared the log-likelihoods of the null and full models (Cf-null and Cf-Mf) to confirm that the relative abundances of the *Methanobacteriaceae* and *Christensenellaceae* families were significantly correlated (χ^2^ test, p-value = 1.78×10^-59^). Furthermore, the model Cf-Mf showed that Mf-tra was significantly positively correlated to Cf-tra (Fig. 1b; type I ANOVA, F-value(339) = 287.03, p-value < 0.0001) and that the interaction term between Mf-tra and age was positively correlated to Cf-tra as well. These results indicate that the relative abundances of the *Christensenellaceae* and *Methanobacteriaceae* families are positively correlated across multiple populations/studies. In addition, although both families are enriched in low-BMI people, they are correlated regardless of a subject’s BMI. Moreover, their association is stronger in older people, suggesting that although elders are less likely to carry as much *Christensenellaceae* as youths, the more *Methanobacteriaceae* they have, the more *Christensenellaceae* they have.

We performed a similar analysis using the abundances of the two most prominent genera belonging to these families (*Christensenella* and *Methanobrevibacter*, models Cg-null and Cg-Mg) and obtained equivalent results. First, the interaction term between Mg-tra and age was positively correlated with Cg-tra (Fig. 1c; type I ANOVA, F-value(339) = 10.19, p-value = 0.0015). Then, by comparing model Cg-null with the full model Cg-Mg, we showed that the relative abundances of the two genera were also correlated (χ^2^ test, p-value = 1.50×10^-57^). Our full model Cg-Mg showed that Mg-tra was significant for predicting Cg-tra while controlling for BMI and age (type I ANOVA, F-value(339) = 274.35, p-value < 0.0001), with the interaction term between Mg-tra and age also positively correlated to Cg-tra. These results indicate that the correlations between the relative abundances of the two families, explained above, hold true at the level of two representative genera. The association between *Methanobrevibacter* and *Christensenella* is stronger in older people regardless of BMI.

A similar analysis at the species level indicated that *C. minuta* and *M. smithii* were the most abundant species of each of their genera, and similarly to the family and genus ranks, their relative abundances across samples were significantly correlated (Fig. 1d and Appendix 2). The less abundant *Christensenella* gut species, *C. massiliensis* and *C*. *timonensis*, also correlated with *Methanobrevibacter smithii* across the 1,821 metagenomes (Appendix 2). *C. minuta* and *C. timonensis* transformed relative abundances were significantly negatively correlated to both BMI and age, while *C. massiliensis* transformed relative abundance was significantly correlated with BMI but not with age. Leaner people are thus enriched in members of the *Christensenellaceae* family, and *C. minuta* and *C. timonensis* are more abundant in younger people than in older people.

### *C. minuta* forms flocs alone and in co-culture with *M. smithii*

To assess the physical and metabolic interaction of two representative species, we used *C. minuta* DSM-22607, previously shown to reduce adiposity in germfree mouse fecal transplant experiments (4), and *M*. *smithii* DSM-861, which is the most abundant and prevalent methanogen in the human gut (31). Confocal and scanning electron imaging of 2-7 day-old cultures revealed that *C. minuta* flocculate in mono- and co-cultures (Fig. 2a,b and Fig. 3a-c, g-j). *M. smithii* is present within the *C. minuta* flocs (Fig. 2d and Fig. 3g-j) but does not aggregate in mono-culture before 7-10 days of culture (data not shown). In contrast, *B. thetaiotaomicron,* used here as a positive control based on previous reports that it supports the growth of *M. smithii* via H_2_ production (25, 26), did not flocculate when grown alone, (Fig. 2c) and, when co-cultured with *M*. *smithii,* displayed very limited aggregation (Fig. 2e, Fig. 3k-n and Fig. S2).

**Fig. 2.**
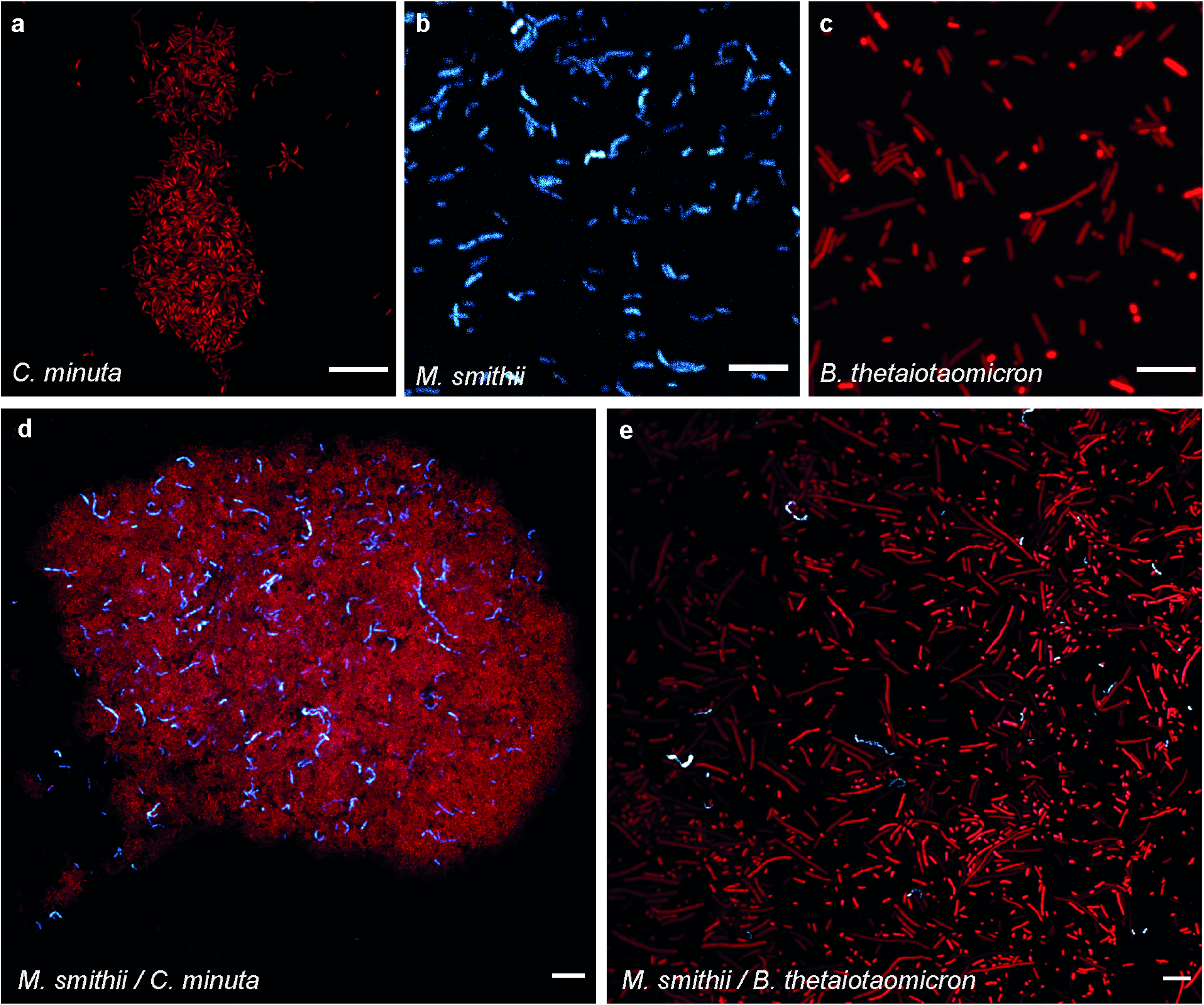
Confocal micrographs of the cultures at 3 days of growth. Confocal micrographs after 3 days of growth. (a) *C. minuta* alone; (b) *M. smithii* alone; (c) *B. thetaiotaomicron* alone; d: *M*. *smithii* and *C. minuta* together; and e: *M. smithii* and *B. thetaiotaomicron* together. SYBR Green I fluorescence (DNA staining) is shown in red and *M. smithii*’s coenzyme F_420_ autofluorescence is shown in blue. Scale bars represent 10 μm. Based on gases production, at 3 days of growth, *B. thetaiotaomicron* was already at stationary phase (explaining the elongated cells, see Fig. S2 for confocal micrographs of *B. thetaiotaomicron* and *M. smithii* at 2 days of growth), *C. minuta* was at the end of the exponential phase and *M. smithii* was still in exponential phase.

**Fig. 3.**
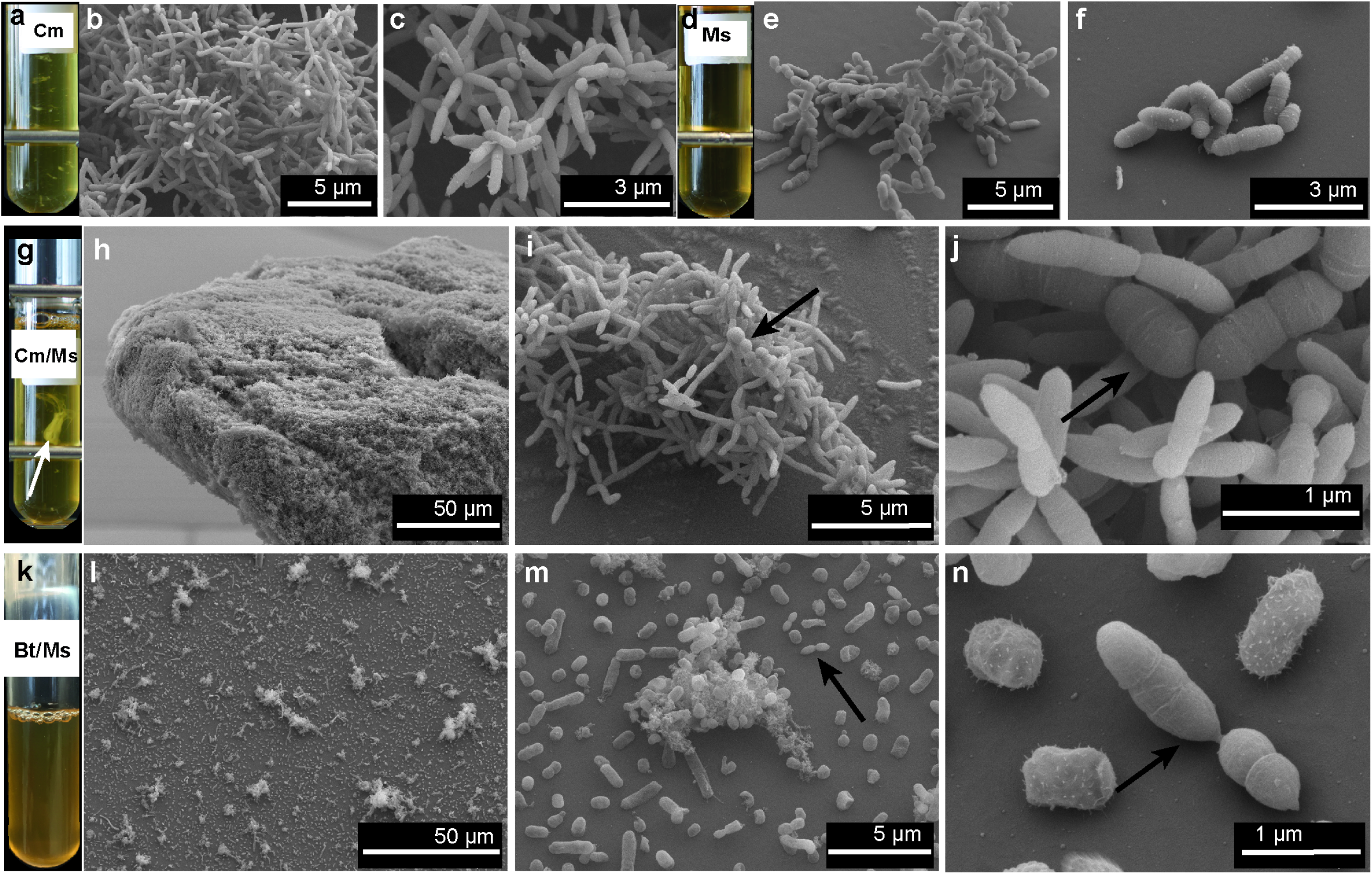
Scanning electron micrographs of the cultures at 3-7 days of growth. (a, d, g and k) Representative Balch tubes of cultures of *C. minuta* (Cm), *M. smithii* (Ms), *C. minuta* and *M. smithii* (Cm/Ms), and *B. thetaiotaomicron* and *M. smithii* (Bt/Ms) after 7 days of growth. In panel g, the floc formed by Cm/Ms is indicated with an arrow. (b-c) Scanning electron micrographs (SEMs) of mono-cultures of *C. minuta* at 5 days of growth; (e-f) SEMs of mono-cultures *M. smithii* at 5 days of growth; (h-j) SEMs of co-cultures of *C. minuta* and *M. smithii* at 7, 5 and 2 days of growth respectively; (l-n) SEMs of co-cultures of *B. thetaiotaomicron* and *M. smithii* at 7 days of growth. Arrows indicate *M. smithii* cells. Metal bars on panels a, d and j are from the tube rack.

### H_2_ and CH_4_ production

After 6 days in mono-culture, *C. minuta* had produced 7 times more H_2_ than *B. thetaiotaomicron* (14.2 ± 1.6 mmol.L^−1^ *vs*. 2.0 ± 0.0 mmol.L^−1^, Fig. 4a and d and Fig. 5a; Wilcoxon rank sum test, p-value = 0.1). As expected, *M. smithii* did not grow in mono-culture when H_2_ was not supplied (80:20 % v/v N_2_:CO_2_ headspace, Fig. 4b). After 6 days, *M. smithii* had produced 9.0 ± 1.0 mmol.L^−1^ of CH_4_ when H_2_ was provided in excess (*i*.*e*., 80:20 % v/v H_2_:CO_2_ atmosphere at 2 bars; Fig. 4b and Fig. 5b).

**Fig. 4.**
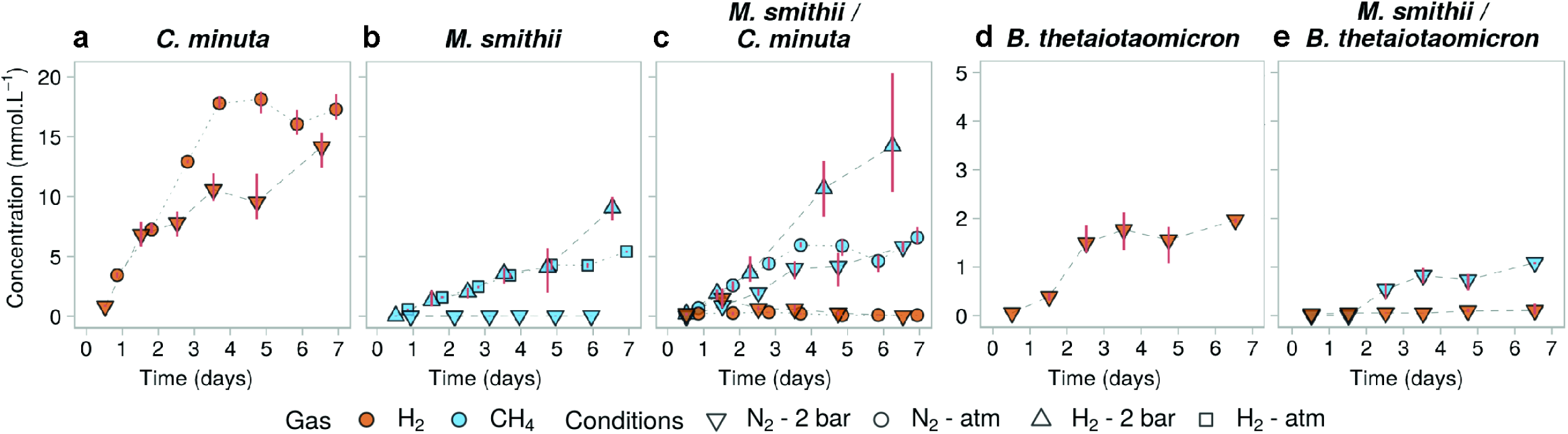
Gas concentrations over time in mono- and co-cultures of *C. minuta*, *B*. *thetaiotaomicron*, and *M. smithii* grown under different conditions. (a-e) H_2_ (orange) and CH_4_ (blue) concentrations in the headspace over time in cultures from batches 1-3 (see Table S1). Points represent the average of the 3 biological replicates for each condition, and red bars join the minimal and maximal values. In conditions where H_2_ was provided in excess (H_2_ - 2 bar and H_2_ - atm, headspace initially composed of 80:20 % H_2_:CO_2_), its concentrations are not shown for scale reasons. Initial concentrations of H_2_ in conditions where it was not provided in the headspace were undetectable (N_2_ - 2 bar and N_2_ - atm, headspace initially composed of 80:20 % N_2_:CO_2_) and stayed null in the mono-cultures of *M. smithii* (not shown). CH_4_ concentrations in the bacterial mono-cultures were undetectable and are not shown as well. Panels a-c share the same y-scale, as do panels d-e.

**Fig. 5.**
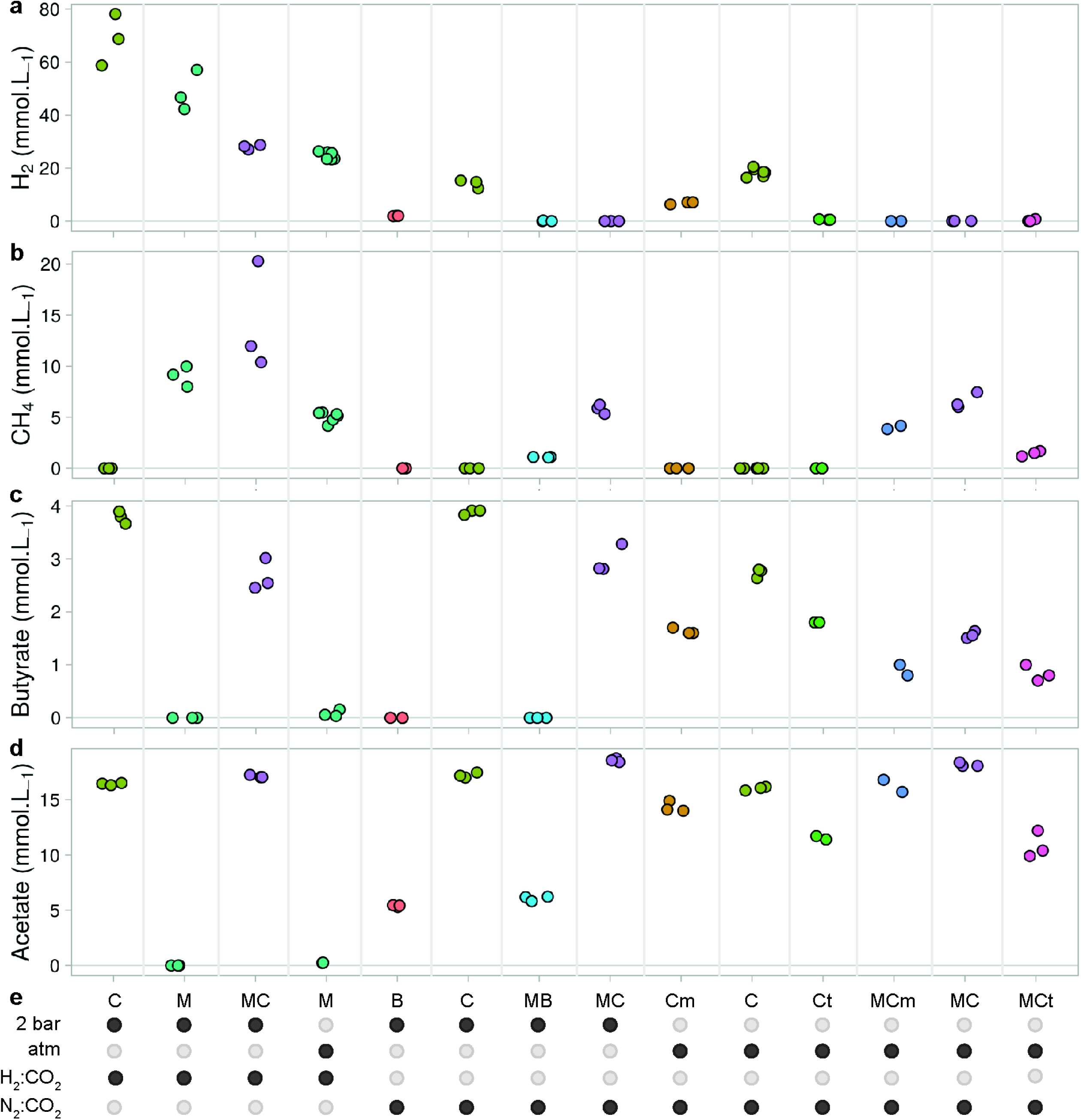
Summary of gases and SCFA produced in mono- and co-cultures of *B. minuta*, *C. timonensis*, *C. massiliensis*, *B. thetaiotaomicron*, and *M. smithii* after 6 days of growth. (a-d) H_2_, CH_4_, butyrate, and acetate production after 6 days of growth in all mono- and co-cultures presented in this study (batches 1-4, Table S1). Points represent the concentration of each biological replicate; (e) Table summarizing the conditions for each culture. The conditions include the gas mixture (H_2_:CO_2_ or N_2_:CO_2_ 80:20 % v/v), the initial pressure (2 bar or atmospheric) and the microorganisms inoculated. C: *C. minuta*. Ct: *C. timonensis*. Cm: *C. massiliensis* (Cm). B: *B. thetaiotaomicron*. M: *M. smithii*. Samples inoculated with the same microorganisms are the same color.

In accordance with the higher levels of H_2_ produced by *C. minuta* compared to *B. thetaiotaomicron*, day-6 CH_4_ concentrations were higher for *M. smithii* co-cultured with *C. minuta* compared to with *B. thetaiotaomicron* (respectively 5.8 ± 0.5 mmol.L^−1^ and 1.1 ± 0.0 mmol.L^−1^; Wilcoxon rank sum test, p-value = 0.1; Fig. 4c and e and Fig. 5b).

For both co-culture conditions, H_2_ concentrations were very low (on average across time points and replicates, H_2_ concentrations were 0.5 ± 0.6 mmol.L^−1^ in co-cultures with *C. minuta*, and 0.1 ± 0.1 mmol.L^−1^ with *B. thetaiotaomicron*), indicating that almost all the H_2_ that had been produced was also consumed (Fig. 4c and e and Fig. 5a).

### Pressure effects on gas production and aggregation

Gas consuming microbes, including hydrogenotrophic methanogens, grow better in a pressurized environment (32–34) due to a higher gas solubility at higher pressure, as described by Henry’s law. We compared CH_4_ production by *M. smithii* in mono-culture and in co-culture with *C. minuta* under 2 different pressures (*i.e*., 2 bar and atmospheric pressure). Similar to the flocculation at 2 bar (Fig. 2d), *C*. *minuta* and *M. smithii* also aggregated at atmospheric pressure (Fig. S3a-b). Accordingly, *C. minuta* supported CH_4_ production by *M*. *smithii* to a similar extent under both pressure conditions (ANOVA followed by Tukey’s post-hoc test, adjusted p-value = 1.0; Fig. 4c, Fig. 5b and, even though the putative H_2_ produced by *C. minuta* (estimated based on the mono-cultures) was much lower than the amount of H_2_ provided in the headspace for *M. smithii* (Fig. 5a).

We next sought to assess if the mixed aggregation of *M. smithii* with *C. minuta* could be disrupted if H_2_ was pressurized in the medium, reducing *M. smithii’*s reliance on *C. minuta* as a H_2_ source. We observed that *M. smithii* aggregated with *C. minuta* (Fig. S3c-d) even though H_2_ was abundant. Total CH_4_ production was higher than in mono-culture under the same headspace, reaching 14.2 ± 5.3 mmol.L^−1^ in co-culture *vs.* 9.0 ± 1.0 mmol.L^−1^ in mono-culture after 6 days (ANOVA followed by Tukey’s post-hoc test, adjusted p-value = 0.1, Fig. 4b and c). This indicates that interspecies H_2_ transfer occurs even when H_2_ is added to the headspace, and leads to greater methanogenesis.

### The short chain fatty acid (SCFA) production of *C*. *minuta* is influenced by the presence of *M*. *smithii*

Regardless of headspace composition and pressure conditions, the only SCFAs detected as produced by *C. minuta* in mono-culture were acetate and butyrate (among 10 short and medium chain fatty acids analyzed, Appendix 1; Fig. 5). To investigate if the consumption of H_2_ by *M. smithii* influenced the SCFA production profile of *C. minuta*, we compared acetate and butyrate concentrations between the co-cultures and *C. minuta*’s mono-cultures under all conditions tested (*i.e*., cultures at 2 bar or atmospheric pressure with an 80:20 % v/v N_2_:CO_2_ or H_2_:CO_2_ headspace, Table S1).

We consistently observed lower butyrate concentrations in all co-cultures compared to mono-cultures (Fig. 6a-c, Fig. 5c; ANOVA, F-value(1) = 161.461 and adjusted p-value = 7.7×10^−8^). For all conditions, butyrate concentrations in co-culture after 6 days were 1.1 ± 0.24 mmol.L^−1^ lower than in mono-cultures (Fig. 6a-c and Table A3). The interaction factor between the mono/co-culture conditions and the growth condition was not significantly correlated to butyrate concentrations (ANOVA, F-value(2) = 0.862, adjusted p-value = 0.4). The observation that butyrate concentrations in co-cultures were lower than in mono-cultures regardless of pressure and headspace composition suggest that the methanogen’s presence shapes the metabolite output of *C. minuta*.

**Fig. 6.**
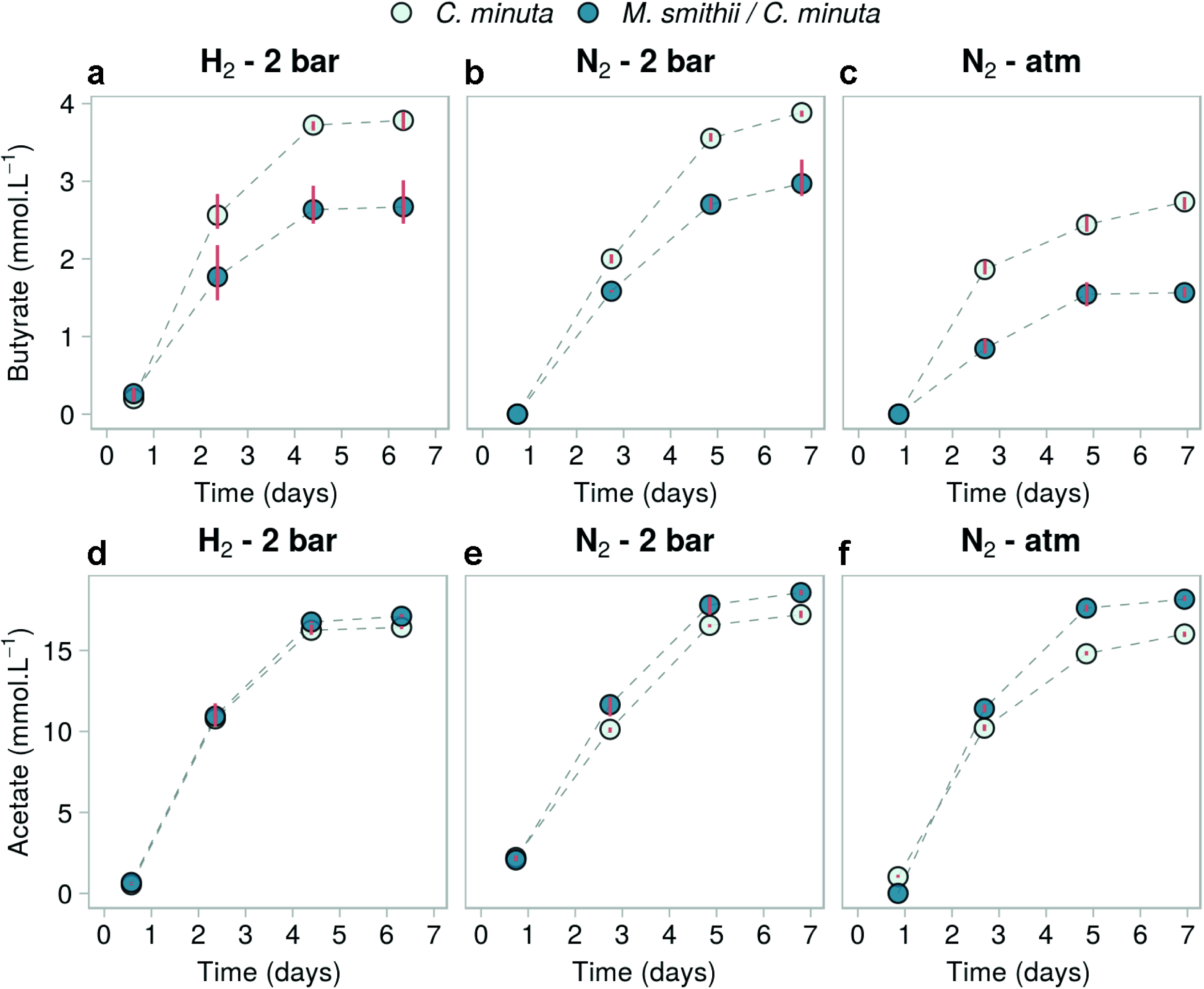
SCFA concentrations over time in mono- and co-cultures of *C. minuta* and *M. smithii* grown under different conditions. Short chain fatty acids over time in cultures from batches 1-3 (see Table S1). (a-c) butyrate concentrations; d-f: acetate concentrations. Only these SCFA were detected among the fatty acids tested (fatty acids from C_1_ to C_8_, iso-valerate and iso-butyrate). Points represent the average of the 3 biological replicates for each condition, and red bars join the minimal and maximal values. Mono-cultures of *M. smithii* are not shown as they did not differ from the blanks (negative controls).

Along with the reduced butyrate production, we also observed slightly but significantly higher acetate production in co-cultures compared to mono-cultures (Fig. 6d-f, Fig. 5d; ANOVA, F-value(1) = 317.41 and adjusted p-value = 3.2×10^−9^). This difference was also observed in three additional batches performed at 2 bars (Fig. S4). The difference in acetate production between mono and co-culture conditions significantly varied with the headspace and pressure conditions (interaction term between the mono- or co-culture and the growth condition was significantly correlated to acetate production; ANOVA, F-value(2) = 29.09 and adjusted p-value = 3.0×10^−5^). The differences in final acetate production (after 6 days) ranged from +0.7 mmol.L^−1^ at 2 bar under an H_2_:CO_2_ (80:20 % v/v) atmosphere to +2.2 mmol.L^−1^ at atmospheric pressure under an N_2_:CO_2_ (80:20 % v/v) atmosphere. Furthermore, we observed in co-culture more CH_4_ than what *M. smithii* could have produced based on the H_2_ production in *C. minuta*’s mono-cultures (Appendix 3). This observation implies that *C. minuta* likely produced a greater amount of H_2_ in the co-cultures along with greater acetate production.

### *C. massiliensis* and *C. timonensis* also support the metabolism of *M. smithii*

We performed similar co-culture experiments of *M. smithii* with *C. massiliensis* and *C. timonensis* at atmospheric pressure. *C. massiliensis* and *C. timonensis* aggregated in mono-culture, and *M. smithii* grew within their flocs in co-culture (Fig. 7). The H_2_ produced by the bacteria in mono-culture after 6 days of growth (6.9 ± 0.5 mmol.L^−1^ for *C. massiliensis* and 0.6 ± 0.1 mmol.L^−1^ for *C. timonensis*, Fig. 8a,d) was lower than the levels produced by *C. minuta* (Fig. 4a). CH_4_ production in the co-cultures reached 4.0 ± 0.2 mmol.L^−1^ with *C. massiliensis* and 1.5 ± 0.3 mmol.L^−1^ with *C*. *timonensis*. These amounts of methane are significantly lower than what we observed for *M. smithii* with *C. minuta* (6.6 ± 0.8 mmol.L^−1^; ANOVA followed by a Tukey’s post-hoc test, adjusted p-values = 6.8×10^-2^ and 1.7×10^−3^ for co-cultures respectively with *C. massiliensis* and *C. timonensis* against *C. minuta*; Fig. 8c,e, and Fig. 4c).

**Fig. 7.**
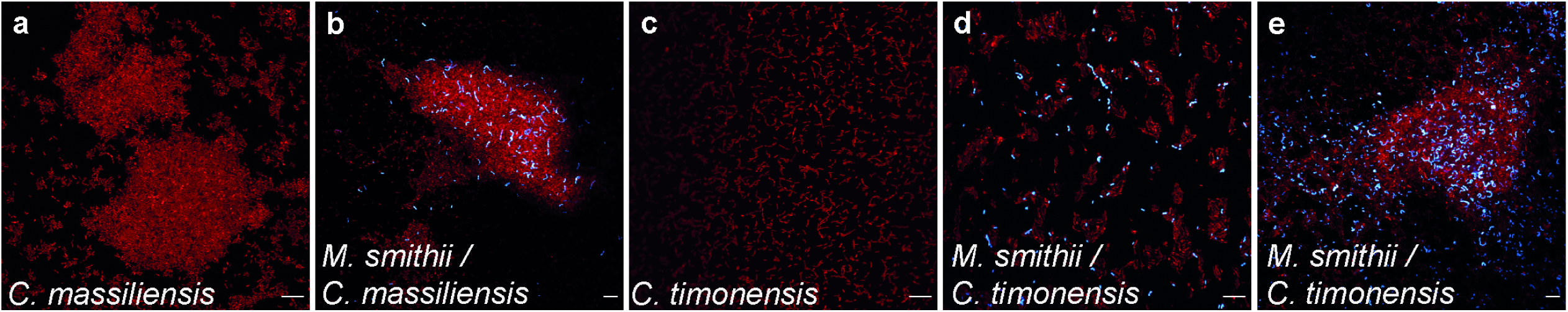
Confocal imaging of *C. massiliensis* and *C. timonensis* in mono- and co-cultures with *M. smithii*. Confocal micrographs after 5 days of growth of (a) *C. massiliensis*, (b) *M. smithii* and *C. massiliensis* in co-culture, (c) *C. timonensis*, d-e: *M*. *smithii* and *C. timonensis* in co-culture. SYBR Green I fluorescence (DNA staining) is shown in red and, *M. smithii*’s coenzyme F420 autofluorescence is shown in blue. Scale bars represent 10 μm.

**Fig. 8.**
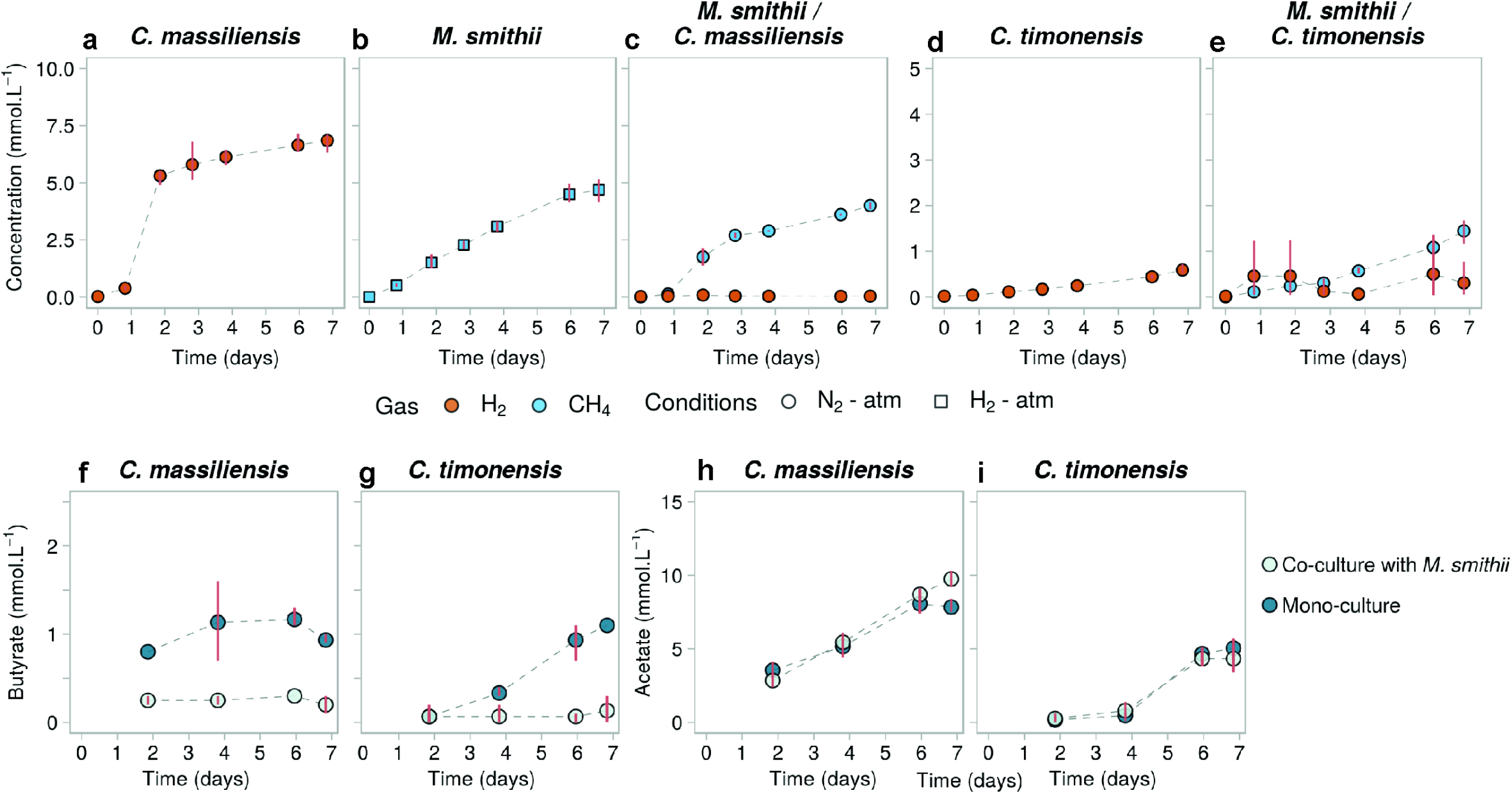
Gas and SCFA concentrations in mono- and co-cultures of *C. massiliensis* and *C. timonensis* with *M. smithii*. (a-e) H_2_ (orange) and CH_4_ (blue) concentrations in the headspace in cultures from batch 4 (see Table S1); (f-g) butyrate and (h-i) acetate concentrations in these cultures. Points represent the average of 3 biological replicates, and red bars join the minimal and maximal values. In the mono-cultures of *M. smithii* (b) where H_2_ was provided in excess (condition H_2_ - atm, headspace initially composed of 80:20 % H_2_:CO_2_), its concentrations are not shown for scale reasons.

We observed less butyrate production in the co-cultures compared to mono-cultures (Wilcoxon rank sum test, p-values = 0.33 for *C. massiliensis* and 0.5 for *C. timonensis*; Fig. 8f,g), with butyrate measured barely above the detection limit in co-cultures. While in mono-cultures, *C. massiliensis* and *C. timonensis* produced 0.93 ± 0.06 mmol.L^-1^ and 1.10 ± 0.00 mmol.L^−1^ of butyrate, respectively; in co-culture with *M. smithii* they produced 0.20 ± 0.14 mmol.L^−1^ and 0.13 ± 0.15 mmol.L^−1^, respectively. Acetate production by *C. massiliensis* was higher in co-culture compared to mono-culture (7.83 ± 0.49 mmol.L^−1^ of acetate produced in mono-culture by day 6 and 9.75 ± 0.78 mmol.L^−1^ produced in co-culture with *M. smithii*), although this difference was not significant (Wilcoxon rank sum test, p-value = 0.2). And in contrast with the co-cultures of *C. minuta* with *M. smithii*, acetate production by *C. timonensis* was not higher in the co-cultures compared to mono-cultures: *C. timonensis* produced 5.05 ± 0.21 mmol.L^−1^ in mono-culture and 4.33 ± 1.21 mmol.L^−1^ in co-culture (Wilcoxon rank sum test, p-value = 0.8; Fig. 8h,i).

## Discussion

The link between the relative abundance of the *Christensenellaceae* and host BMI now stands as one of the most reproducible associations described between the gut microbiome and obesity (4–15). Here, we confirm in a meta-analysis of metagenomes across 10 populations, the previously observed association between leanness and the *Christensenellaceae* family (4, 20–22). We could also show that *Christensenella* genus and *Christensenella* spp. also correlated with leanness. Similarly, we observed correlations between leanness and the *Methanobacteriaceae* family, the *Methanobrevibacter* genus and *M. smithii*. These methanogens were positively correlated with members of the *Christensenellaceae* family. The relative abundances of the *Christensenellaceae* were higher in young people, whereas conversely, *Methanobacteriaceae* was enriched in older people. Despite these opposite patterns, the two families correlate with each other regardless of age and BMI.

We selected the two most prominent members of the two families, *C. minuta* and *M. smithii*, to ask if physical and metabolic interactions could underlie these positive associations. *C. minuta* produced copious amounts of H_2_ during fermentation. In co-culture with *C. minuta*, *M. smithii* produced comparable amounts of CH_4_ as in mono-culture with an excess of H_2_, indicating that *C. minuta* can efficiently support the growth of *M. smithii* via interspecies H_2_ transfer. *C. minuta* formed flocs visible by eye, and *M. smithii* grew within these flocs.

*M. smithii* would likely benefit by associating with the flocs formed by *C. minuta* through better access to H_2_. Interspecies metabolite transfer corresponds to the diffusion of a metabolite (*e.g*., H_2_) from the producer (*e*.*g*., *C. minuta*) to the consumer (*e*.*g*., *M. smithii*). As described by Fick’s law of diffusion, the flux of a metabolite between two microorganisms is directly proportional to the concentration gradient and inversely proportional to the distance, such that the closer the microorganisms are, the better the H_2_ transfer (35, 36). Thus, within the flocs the H_2_ interspecies transfer would be more efficient, to the benefit of *M. smithii*. In accord, we observed greater methane production under excess H_2_ when *C. minuta* was present.

When grown in co-culture, *M*. *smithii* influenced the metabolism of *C. minuta*. The presence of the methanogen inhibited the production of butyrate while enhancing acetate production by *C. minuta* under all growth conditions, on average among all experimental batches. This observation suggests that H_2_ consumption by *M. smithii* decreased the P_H2_ within the floc enough to favor acetate production (37). The consumption of H_2_ causes the cell to produce more oxidized fermentation products such as acetate (38–41), and the interspecies H_2_ transfer leads to greater CH_4_ production.

Both the methane production and the co-flocculation were far more pronounced when *M. smithii* was grown with *C. minuta* compared to with *B. thetaiotaomicron*. *B*. *thetaiotaomicron* has previously been shown to support the growth of *M*. *smithii* in co-culture (25, 26). *B*. *thetaiotaomicron* barely aggregated, in contrast to *C. minuta*’s very large (visible to the naked eye) flocs. When grown together, *B*. *thetaiotaomicron* and *M. smithii* showed very poor aggregation. Moreover, acetate was the only SCFA detected in mono-cultures of *B. thetaiotaomicron*, and its production was less affected by the methanogen compared to *C. minuta*. Methane produced by *M. smithii* in co-culture with *B. thetaiotaomicron* was one fifth of that produced with *C. minuta*, possibly as a result of the smaller amount of H_2_ produced and the reduced contact between cells. Given that *M. smithii* does not co-occur with *B. thetaiotaomicron* in human microbiome datasets, this is another indication that co-occurrence patterns may point to metabolic interactions.

*C. massiliensis* and *C. timonensis* also produced H_2_, acetate and butyrate, and also flocculated in mono-culture. *C. massiliensis* and *C. timonensis* supported methane production by *M. smithii,* which grew within the bacterial flocs. However, *M. smithii* also grew outside the flocs when co-cultured with these two species, which we did not observed in the co-cultures with *C. minuta*. And although *M. smithii* also influenced fermentation of *C. massiliensis* and *C. timonensis*, the overall changes in SCFA production in co-culture were different from what we observed with *C. minuta*: butyrate production was almost undetectable, while acetate production was not significantly affected.

These results suggest that the interaction between *M. smithii* and *C. minuta* leads to higher methane production compared to *B. thetaiotaomicron* and to other species of the *Christensenellaceae*, possibly due to the higher levels of interspecies H_2_ transfer. Nevertheless, *C. massiliensis* and *C. timonensis* did support CH_4_ production better compared to *B. thetaiotaomicron*. The higher H_2_ production of *C. massiliensis* compared to *B. thetaiotaomicron* might explain this. In the case of *C. timonensis*, although it produced half of the H_2_ produced by *B. thetaiotaomicron* in mono-culture, *M. smithii* produced more CH_4_ in co-culture with *C. timonensis* than with *B. thetaiotaomicron*. This suggests that, similar to its effect on *C. minuta*, *M. smithii* also triggered the production of H_2_ by *C. timonensis*.

Altogether, our work demonstrates that members of the *Christensenellaceae* act as a H_2_ source to methanogens, and this process is enhanced via close physical proximity. Such interactions also likely underlie the co-occurrence patterns of the *Christensenellaceae* with other members of the microbiome. Many of these families lack cultured representatives, such as the *Firmicutes* unclassified SHA-98, *Tenericutes* unclassified RF39 and unclassified ML615J-28 (4). Based on our results, cultivation of these elusive members of the microbiome may require H_2_ (or the provision of another metabolite that *C. minuta* produces when H_2_ is being consumed). Despite their very low abundance in the human gut, members of the *Christensenellaceae* may shape the composition of the gut microbiome by favoring the colonization and persistence of certain hydrogenotrophs, and by supplying other butyrate producers with acetate (42).

Here, we confirmed an association of *M. smithii* and leanness based on metagenomes from 10 studies. In contrast, some studies have reported an association between *M. smithii* and obesity (2, 43). In this scenario, H_2_ uptake by *M. smithii* would promote the breakdown of non-digestible carbon sources by fermenters, such as acetogens, thereby increasing the amount of acetate or other SCFA that can be absorbed and utilized by the host and promoting fat storage (2, 44). In contrast, and consistent with our results, *M. smithii* has also been repeatedly associated with anorexia and leanness (4, 45–48). In this case, the production of CH_4_ would decrease the amount of energy available for the host via carbon loss, as has been observed in livestock (49–52). Thus our observation, that the presence of *M. smithii* directs the metabolic output of the *C. minuta* towards greater H_2_ availability for methanogenesis, via increased acetate production, is consistent with their association with a lean phenotype. To assess quantitatively how the presence and activity of these microbes impact host physiology will require careful modeling of energy flow *in-vivo*.

## Materials and methods

### Metagenome data generation

We generated 141 metagenomes from fecal samples obtained as part of a previous study (53) (Supplementary Table S2). Metagenomic libraries were prepared as described in Appendix 1, Additional methods.

### Data from public databases

We constructed a metagenome sequence collection from: i) the newly generated data (above) to complement the 146 metagenomes previously reported in Poole et al., 2019 (53); and ii) publicly available shotgun-metagenome sequences from stool samples included in the curatedMetagenomicData package of Bioconductor (54) for which BMI information was provided. For the latter, we restricted our analyses to individuals for which the following information was available: gender, age, country of origin, and BMI. Individuals with *Schistosoma* (n = 4), or Wilson’s disease (n = 2), were excluded from the analysis, as were samples from two pregnant women. In all, 1,534 samples from 9 studies were downloaded from the sequence read archive (SRA) and further processed (Table S3) for a total or 1,821 samples with at least one million sequence pairs per sample.

### Data processing

A detailed description of the processing of the raw sequences is given in Appendix 1. To obtain a taxonomic profile of the metagenome samples, we built a custom genomes database (55) for Kraken v2.0.7 (56) and Bracken v2.2 (57) using the representative genomes from the Progenomes database (as available on August 24 2018) (58), to which we added genome sequences of *C. minuta* (GenBank assembly accession: GCA_001652705.1), *C. massiliensis* (GCA_900155415.1), and *C. timonensis* (GCA_900087015.1). Reads were classified using Kraken2 and a Bayesian re-estimation of the species-level abundance of each sample was then performed using Bracken2. We obtained complete taxonomic annotations from NCBI taxIDs with TaxonKit v0.2.4 (https://bioinf.shenwei.me/taxonkit/). The detection limit for the relative abundances in samples was 10^-3^ %; in consequence, all relative abundances below this threshold were equal to 0.

### Meta-analysis of human gut metagenomes

Linear mixed models (R package nlme) were used to evaluate the correlation between the relative abundances of taxa while correcting for the structure of the population; the study of origin was set as a random effect. In some datasets, individuals were sampled multiple times in which case the individual effect was nested inside the dataset effect. Relative abundances were transformed using the Tukey’s ladder of powers transformation (59), and are designated with the suffix ‘-tra’ (*e.g.*, the transformed relative abundance of the family *Christensenellaceae* is Cf-tra). Covariates in null models were selected using a backward feature selection approach based on a type II ANOVA (*i.e*., by including all covariates and removing the non-significant ones step-by-step until all remaining variables were significant, Appendix 2). We made 4 null models predicting the transformed relative abundance of the family *Christensenellaceae* (Cf-null), the genus *Christensenella* (Cg-null), the family *Methanobacteriaceae* (Mf-null) and the genus *Methanobrevibacter* (Mg-null). To evaluate the correlation between taxa, we made model Cf-Mf by adding Mf-tra and its interaction with age to the covariates of Cf-null.

Reciprocally, we made model Cg-Mg by adding Mg-tra and its interaction with age to the covariates of Cg-null. The same approach was performed at the species level and it is described in Appendix 2.

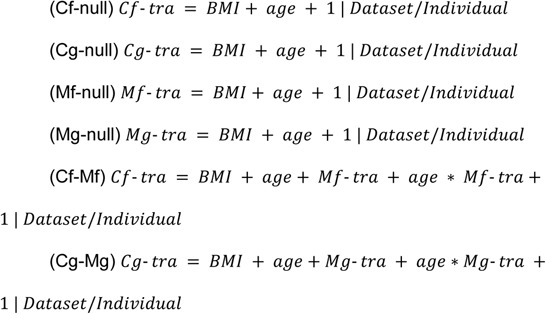

We used the likelihood ratio test to compare the nested models via the χ^2^ distribution (*i.e.* Cf-null *vs.* Cf-Mf and Cg-null *vs.* Cg-Mg). To characterize the correlation of Cf-tra with Mf-tra, and Cg-tra with Mg-tra, after correcting for BMI and age, we used a type I ANOVA to evaluate the importance of the variables in the order they appear in Cf-Mf and Cg-Mg. The F-value, degree of freedom and p-value are reported for each variable. All analyses were performed using R (60).

### Culturing of methanogens and bacteria

We obtained *M. smithii* DSM-861, *C. minuta* DSM-22607, *C. massiliensis* DSM 102344, *C. timonensis* DSM 102800, and *B. thetaiotaomicron* VPI-5482 from the German Collection of Microorganisms and Cell Cultures (DSMZ; Braunschweig, Germany). Each culture was thawed and inoculated into Brain Heart Infusion (BHI) medium (Carl Roth, Karlsruhe, Germany) supplemented with yeast extract (5 g/L), reduced with L-Cysteine-HCl (0.5 g/L) and Ti-NTA III (0.3 mM), and buffered with sodium bicarbonate (42 mM, pH 7, adjusted with HCl 6M). 10 mL cultures were grown at 37°C without shaking in Balch tubes (total volume of 28 mL) under a headspace of N_2_:CO_2_ (80:20% v/v) in the case of the bacteria, and H_2_:CO_2_ (80:20% v/v, pressure adjusted to 2 bar) for *M. smithii*. When initial cultures reached exponential growth, and before floc formation, they were transferred into fresh medium and these transfers were used as inocula for the experiments described below.

### Co-culture conditions

*M. smithii* was co-cultured with *C. minuta*, *B. thetaiotaomicron*, *C. massiliensis*, or *C*. *timonensis*, and in parallel, each microorganism was grown in mono-culture (Table S1). Prior to inoculation, one-day old cultures of bacterial species, or 4-day old cultures of *M. smithii*, were adjusted to an OD_600_ of 0.01 with sterile medium. For the co-cultures, 0.5 mL of each adjusted culture was inoculated into 9 mL of fresh medium. For the mono-cultures, 0.5 mL of the adjusted culture and 0.5 mL of sterile medium were combined as inoculum. For negative controls, sterile medium was transferred as a mock inoculum. Headspaces were exchanged with 80:20 % (v/v) of N_2_:CO_2_ or H_2_:CO_2_ and pressurized at 2 bar or atmospheric pressure (Table S1). Each batch of experiments was carried out once with 3 biological replicates per culture conditions (Table S1).

### Imaging

For confocal microscopy, SYBR^®□^ Green I staining was performed as previously described (61) with the modifications described in Appendix 1. Imaging by confocal microscopy (LSM 780 NLO, Zeiss) was used to detect the autofluorescence emission of coenzyme F_420_ of *M. smithii* and the emission of SYBR Green I (Appendix 1). Images were acquired with the ZEN Black 2.3 SP1 software and processed with FIJI (62). Micrographs are representative of all replicate cultures within each experimental batch. The preparation of the samples for scanning electron microscopy is described in Appendix 1. Cells were examined with a field emission scanning electron microscope (Regulus 8230, Hitachi High Technologies, Tokyo, JPN) at an accelerating voltage of 10 kV.

### Gas and SCFA measurements

Headspace concentrations of H_2_, CO_2_, and CH_4_ were measured with a gas chromatograph (GC) (SRI 8610C; SRI Instruments, Torrence, USA) equipped with a packed column at 42°C (0.3-m HaySep-D packed Teflon; Restek, Bellefonte, USA), a thermal conductivity detector (TCD) at 111°C, and a flame ionization (FID) detector. The gas production and consumption were estimated from the total pressure in the vials (ECO2 manometer; Keller, Jestetten, Germany) and the gas concentrations in the headspace using the ideal gas equation. The concentrations are given in mmol of gas in the headspace per liter of culture.

SCFA measurements were performed with liquid samples (0.5 mL) filtered through 0.2 μm pore size polyvinylidene fluoride filters (Carl Roth, GmbH, Karlsruhe, GER). SCFA concentrations were measured with a CBM-20A high performance liquid chromatography (HPLC) system equipped with an Aminex HPX-87P column (300 x 7.8 mm, BioRad, California, USA), maintained at 60 °C, and a refractive index detector. A sulfuric acid solution (5 mM) was used as eluent at a flow rate of 0.6 mL/min (**∼**40 bar column pressure). Calibration curves for acetate and butyrate were prepared from 1.25 to 50 mM using acetic acid and butyric acid, respectively (Merck KGaA, Darmstadt, Germany). No other fatty acids were detected (Appendix 1). The SCFA concentrations were estimated with the Shimadzu LabSolutions software.

### Statistical analyses

We used Wilcoxon rank sum tests to compare gas production between cultures after 6 days of growth. We performed ANOVA tests when more than one culture condition (*i.e*., headspace composition and pressure, Table S1) was included in the comparison. The conditions in the ANOVA tests (*i.e*., headspace composition and pressure, in mono- or co-culture) were evaluated to explain the variance of CH_4_ production after 6 days of growth. A Tukey’s post-hoc test was then performed to discriminate between the effects of the different conditions. SCFA concentrations were compared using a two-way ANOVA where the culture conditions (*i.e*., headspace composition and pressure, Table S1) and the sample (mono- and co-culture) were evaluated to explain the variance of butyrate and acetate concentrations after 6 days of growth. p-values were adjusted using the Benjamini-Hochberg method. A Tukey’s post-hoc test was performed to discriminate between the effects of the different conditions. All statistical analyses were done in R using the stats R package.

### Data and code availability

The metagenomic sequence data generated during this study have been deposited in the European Nucleotide Archive with accession IDs PRJEB34191 (http://www.ebi.ac.uk/ena/data/view/PRJEB34191). The jupyter notebooks and associated data are available at https://github.com/Albabune/Ruaud_EsquivelElizondo.

## Acknowledgments

We are grateful to Monika Temovska, Sophie Maisch and Iris Holdermann for valuable help. We also thank Daren Heavens for the metagenome library preparation protocol. This work was supported by the Max Planck Society and the Humboldt Foundation.

## Appendixes

## Appendix 1. Additional methods

### Metagenomic libraries preparation

Metagenomic libraries were prepared using 1 ng of DNA input per sample (extracted with the MagAttract PowerSoil DNA kit, Qiagen) as previously described (63). Fragment sizes were restricted to 400 - 700 bp using BluePippin (Sage Science), and samples were pooled at equimolar concentrations before being run on an Illumina HiSeq3000 with 2×150 bp paired end sequencing, resulting in sequencing depths of 3.0 ± 2.1 Gb (median ± standard deviation).

### Raw Data processing

Raw sequences were first validated using fqtools v.2.0 (64) and de-duplicated with the “clumpify” command of bbtools v37.78 (https://jgi.doe.gov/data-and-tools/bbtools/). We trimmed adapters and performed read quality control using skewer v0.2.2 (65) and the “bbduk” command of bbtools. We used the “bbmap” command of bbtools to filter human genome reads that mapped to the hg19 assembly. Finally, we generated QC reports for all reads with fastqc v0.11.7 (https://github.com/s-andrews/FastQC) and multiQC v1.5a (66).

### Confocal imaging, equipment, and settings

For confocal microscopy, SYBR^®️^ Green I staining was performed as previously described (61) with the following modifications: 0.5 mL of culture were sampled and pelleted by centrifugation for 6 min at 6,000 xg (Benchtop centrifuge, Eppendorf, Hamburg, Germany) and pellets were resuspended in a solution containing 744 μL 1x PBS, 16 μL 25x SYBR^®️^ Green I (Sigma-Aldrich, Merck, Germany) and 40 μL 70% v/v ethanol. Samples were pelleted and resuspended before imagining in 100 μL 1x PBS, of which 5 μL were immobilized on 50 μL solid agar (1.5% noble agar in distilled water) (67). Imaging was performed with a confocal microscope (LSM 780 NLO, Zeiss) using oil and water objectives (40x and 63x). A DPSS laser at 405 nm was used to excite the F_420_ enzyme of *M. smithii.* Autofluorescence emission was collected on a 32 channel GaAsP array from 455 to 499 nm. A transmitted light detector (T-PMT) was used to collect the whole light spectrum to create a bright field image. On a second track, an Argon laser at 488 nm was used to excite SYBR^®□^ Green I and its emission was collected from 508 to 588 nm with the 32 channel GaAsP array as well. Images were acquired with a time and space resolution of 2048×2048x(1 to 12)x (xyzt) and pixel dimensions of 0.1038×0.1038 μm for the images taken with the x40 oil objective and pixel dimensions of 0.0659×0.0659 μm for the images taken with the x63 oil objective. The bit depth was 16-bit. Acquisition was performed at 20 °C.

### Processing of the confocal images

FIJI (62) was used to process the confocal micrographs. Contrast and brightness adjustment were applied to the whole image. Due to the thickness of the aggregates of *Christensenella minuta*, the SYBR^®□^ Green I fluorescence intensity was varied with different focal planes. We used a gamma transformation (with gamma = 0.50) to homogenize the fluorescence intensity. The exact same transformation was applied to all samples, even though there were no aggregates, for consistency purposes. Similarly, we applied a gamma transformation to the F_420_ autofluorescent channel to decrease the low fluorescence coming from SYBR^®□^ Green I (gamma = 1.20 to 1.50). As their excitation and emission spectra overlap, there was a low fluorescence intensity of the SYBR^®□^ Green I on the F_420_ autofluorescent channel. The lookup tables (LUT) were Cyan Hot for the F_420_ autofluorescence and red (linear LUT, covering the full range of the data) for the SYBR^®□^ Green I fluorescence.

### Preparation of samples for scanning electron microscopy

Pellets were washed 3-5 times with 1x PBS and then fixed with a 2.5% v/v glutaraldehyde solution in 1x PBS for 1-2 h at room temperature and post-fixed with 1% w/v osmium tetroxide for 1h on ice. Samples were dehydrated in a graded ethanol series followed by drying with CO_2_ in a Polaron critical point dryer (Quorum Technologies, East Sussex, UK). Finally, cells were sputter coated with a 5 nm thick layer of platinum (CCU-010 Compact coating unit, Safematic GmbH, Bad Ragaz, SWI).

### Screening of the short and medium chain fatty acids produced

Before carrying out the experiments presented in the main text, we used gas chromatography (GC) to determine which fatty acids were produced by the cultures and if the corresponding peaks were present in BHI. For this screening, the external standards included equimolar mixtures of acetate, propionate, iso-butyrate, butyrate, iso-valerate, valerate, iso-caproate, caproate, heptanoate, and caprylate, from 0.2 to 7 mM. Measurements were performed with a 7890B GC system (Agilent Technologies Inc., Santa Clara, USA) equipped with a capillary column (DB-Fatwax UI 30 m x 0.25 m; Agilent Technologies) and an FID detector with a ramp temperature program (initial temperature of 80 °C for 0.5 min, then 20 °C per min up to 180°, and final temperature of 180 °C for 1 min). The injection and detector temperatures were 250 and 275 °C, respectively. Samples were prepared as for HPLC (Methods in the main text) with the addition of an internal standard (Ethyl-butyric acid) and acidification (to pH 2) with 50% formic acid. Data were acquired and analysed with the Agilent OpenLAB CDS software.

Only acetate and butyrate were detected in the mono- and co-cultures, and none of the other short and medium chain fatty acids used as standards were detected. As formate was used to acidify samples for the GC measurements, to assess if it was a main product in the cultures, its concentration was measured by HPLC. We also looked for ethanol using HPLC but similar to formate, it was not detected in any of the cultures.Thus, for the experiments in the main text, only acetate and butyrate were quantified via HPLC. BHI medium showed peaks corresponding to 0.33 mM formate and 6 mM of acetate, which were subtracted from the reported concentrations of the cultures.

## Appendix 2. Additional statistics

### Variable selection for the null model

To construct the null model, we tested the effect of the following covariates with a marginal ANOVA: sequencing depth, gender, country, BMI, and age. The sequencing depth was not significant (p-value = 0.73) and was subsequently removed. BMI and age were correlated with the *Chistensenellaceae* abundance (p-value = 0.0002 for the correlation with BMI and 0.02 for the correlation with age).

As *Methanobrevibacter smithii* has been associated with age (23) and BMI (4, 43, 45–48, 68, 69), we first added the interaction factors to the models: the interaction factors were not significant with BMI (p-values = 0.11 and 0.07, for both *Methanobrevibacter* and *Methanobacteriaceae*, respectively). However, the interactions with age were significant (p-values = 0.004 and 0.002, for both *Methanobrevibacter* and Methanobacteriaceae, respectively) and therefore, both variables were kept in the models.

### Statistical analysis at the species rank

Similar to the analysis at family and genus levels presented in the main text, we performed an analysis at the species level between *Christensenella minuta* and *Methanobrevibacter smithii*, the most abundant and prevalent species of their genera. We also studied the correlation of the other two known species of *Christensenella*, *i.e*., *C. massiliensis* and *C. timonenesis*, with *M. smithii*. *M. smithii* was detected in 78.7 % of the samples with a mean relative abundance of 0.53 % (*M. oralis* was the only other *Methanobrevibacter* detected, with a prevalence of 42.8 % and a mean relative abundance of 3.07×10^-3^ %). *C. minuta* had an averaged relative abundance of 0.05% in the 99.7% samples where it was present. *C. timonensis* and *C. massiliensis* had respectively, prevalences of 95.11 % and 98.57 % and mean relative abundances of 6.49×10^-3^ % and 0.02 %.

*M. smithii* was significantly positively correlated with age (type II ANOVA, F-value = 13.22 and p-value = 2.86×10^-4^) and negatively correlated with BMI (type II ANOVA, F-value = 4.13 and p-value = 0.04). The association between *M. smithii* and leanness was not as strong as for its family and genus levels, meaning that other *Methanobacteriaceae* members must contribute to the strength of the association.

Consistently with the analyses at the family and genus levels, *Christensenella minuta* and *Methanobrevibacter smithii* were significantly correlated (χ^2^ test, p-value = 1.05×10^-34^) and the effect of *M. smithii* was significant (type I ANOVA, p-value < 0.0001, F-value = 147.82). Moreover, *C*. *minuta*’s relative abundance correlated with both age and BMI (type I ANOVA, p-values = 0.0071 and 0.0011, F-values = 7.28 and 10.91, respectively), as well as with the interaction term between *M. smithii* and age (type I ANOVA, p-value < 0.0001, F-value = 17.99).

*Christensenella timonensis* and *M. smithii* were correlated (χ^2^ test, p-value = 6.12×10^-98^; type I ANOVA, p-value < 0.0001, F-value = 482.42). And, similar to *C. minuta*, the relative abundance of *C. timonensis* correlated with both age and BMI (type I ANOVA, p-values = 0.0012 and 0.0001, F-values = 10.59 and 16.61, respectively), as well as with the interaction term between *M. smithii* and age (type I ANOVA, p-value < 0.0001, F-value = 35.50).

The relative abundance of *Christensenella massiliensis* correlated with BMI (type I ANOVA, p-value = 0.0028 and F-value = 9.0804) but not with age (p-value > 0.5) and so, we did not correct for age in the null model nor in the model including *Methanobrevibacter smithii*’s transformed relative abundance. *C. massiliensis* and *M. smithii* were also correlated (χ^2^ test, p-value = 1.31×10^-61^; type I ANOVA, p-value < 0.0001, F-value = 310.51), but the interaction term between *M. smithii* and age was not significantly correlated to the bacterium’s abundance. *C. massiliensis* is thus the only *Christensenella* spp. for which the correlation with the methanogen abundance is not a function of the age of the carrier.

## Appendix 3. Comparison of the expected (theoretical) *vs*. the measured methane production in co-cultures

We used the stoichiometry of hydrogenotrophic methanogenesis (CO_2_ + 4 H_2_ = CH_4_ + 2 H_2_O) to calculate the amount of CH_4_ that could be produced from the estimated amount of H_2_ consumed in each sample. For this, we used the mono-cultures of bacteria as references and assumed equal H_2_ production in co-culture as in mono-culture. We estimated the H_2_ consumed after 6 days for each replicate as the difference between the averaged H_2_ concentrations in mono-cultures and the concentration measured in co-culture (*i*.*e*., unconsumed H_2_). The estimated H_2_ consumed was then divided by 4 in order to obtain the theoretical amount of CH_4_ that could be produced via hydrogenotrophic methanogenesis.

**Table A3.** Analysis of the origin of the high methane produced in co-culture based on the changes in metabolism of C. minuta. CH_4_ produced in co-culture was higher than the theoretical amount of CH_4_ that could be generated from H_2_ assuming that *C. minuta* produced the same amount of H_2_ in both mono- and co-cultures. The additional CH_4_ observed could originate from the shift in metabolism from butyrate to acetate production along with H_2_ by *C. minuta* in co-culture. The average concentration among the triplicates after 6 days of growth is given with the standard deviation (SD).

| Condition | $\text{H}_2:\text{CO}_2$ - 2 bar <sup>a</sup> | | $\text{N}_2:\text{CO}_2$ - atm | | $\text{N}_2:\text{CO}_2$ - 2 bar | |
| --- | --- | --- | --- | --- | --- | --- |
|  | Average | SD | Average | SD | Average | SD |
| $\text{H}_2$ produced in mono-culture ($\text{mmol.L}^{-1}$) | 18.71 | 9.71 | 17.28 | 1.12 | 14.15 | 1.56 |
| $\text{H}_2$ not consumed in co-culture <sup>b</sup> ($\text{mmol.L}^{-1}$) | -21.81 | 0.87 | 0.08 | 0.01 | 0.03 | 0.00 |
| Theoretical $\text{CH}_4$ produced based on $\text{H}_2$ produced in mono-culture ($\text{mmol.L}^{-1}$ <sup>c</sup>) | 10.13 | 0.22 | 4.30 | 0.00 | 3.53 | 0.00 |
| Observed CH<sub>4</sub> produced in co-culture (mmol.L<sup>-1</sup>) | 14.21 | 5.33 | 6.57 | 0.77 | 5.81 | 0.45 |
| Difference between observed and theoretical CH<sub>4</sub> (mmol.L<sup>-1</sup>) | 04.08 | 5.50 | 2.27 | 0.77 | 2.28 | 0.45 |
| Butyrate difference between co- and mono-culture (mmol.L<sup>-1</sup>) | -1.11 | 0.30 | -1.21 | 0.04 | -0.91 | 0.27 |
| Acetate difference between co- and mono-culture (mmol.L<sup>-1</sup>) | 0.68 | 0.10 | 2.20 | 0.22 | 1.36 | 0.17 |

<sup>a</sup> For the experiments grown under an H<sub>2</sub>:CO<sub>2</sub> (80:20 %) atmosphere, the average H<sub>2</sub> concentration
measured in the negative controls after 6 days (sampled as many times as the cultures) was subtracted
from the concentration measured in the cultures.
<sup>b</sup> Average of the concentration of H<sub>2</sub> in co-cultures to which the average of H<sub>2</sub> concentration in mono-
culture of *C. minuta* was subtracted.
<sup>c</sup> This amount is calculated based on the stoichiometry of the hydrogenotrophic methanogenesis reaction:
$4 \text{ H}_2 + \text{CO}_2 = \text{CH}_4 + 2 \text{ H}_2\text{O}.$

---

**Fig. S1. Abundances of the *Methanobacteriaceae* and *Christensenellaceae* families across studies.** (a-j) Transformed relative abundances of *Christensenellaceae* (Cf-tra) and *Methanobacteriaceae* (Mf-tra) across 1,821 samples from 10 countries and generated from 10 independent studies. The data generated for this study are grouped with the first time series published in Poole et al., 2019. The gap between 0 and ∼0.2 is due to the detection limit of the sequencing method; the minimal relative abundance is 10^-3^ %. Hence, 0 indicates the microorganism was not detected, which introduces a gap after transformation of the data.

**Fig. S2. Confocal imaging of co-cultures of *B. thetaiotaomicron* and *M. smithii* at different time points.** (a-b) cells at day 2, when *B. thetaiotaomicron* enters stationary phase (see Fig. 4d). (c-d) at day 7, the end of the experiment, when maximal CH_4_ concentrations were observed both in mono-cultures of *M. smithii* and in co-cultures with *B. thetaiotaomicron* (Fig. 4b,e). In exponential phase, *B. thetaiotaomicron* cells are rod-shaped (a); while during stationary phase they suffer stress, leading to elongated cells (c). The bright fields (a and c) and *M. smithii*’s co-enzyme F_420_ (b and d) channels are displayed. Scale bars represent 10 μm.

**Fig. S3. *C. minuta* and *M. smithii* also aggregate at atmospheric pressure and even when there is excess H_2_ in the medium.** Confocal imaging of *C. minuta* and *M. smithii* at 3 days of growth. (a-b) co-culture grown at atmospheric pressure; (c-d) co-culture grown under a pressurized H_2_:CO_2_ atmosphere. The bright fields (a and c) and *M. smithii*‘s co-enzyme F_420_ (b and d) channels are displayed. Scale bars represent 10 μm.

**Fig. S4. Additional batches.** H_2_, CH_4_, acetate, and butyrate concentrations in mono- and co-cultures of *M. smithii* and *C. minuta* grown at 2 bars, as described in the main text. The SCFA of batch S1 were measured by Gas Chromatography instead of High-Performance Liquid Chromatography. The points represent the average of 2 to 3 biological cultures, and red bars join the minimal and maximal values.

## Tables

**Table S1.** Total pressure, headspace composition, and culture inocula for each batch of experiments described in the main text.

**Table S2.** Samples from the Poole et al. (2019) study. Additional data were generated from time points that had not been sequenced for the Poole et al. study. For each individual and each time point, the color indicates whether the sample was prepared and sequenced as previously (orange) or as described in the Methods (blue). If the sample was failed sequencing or was otherwise missing, the color is white.

**Table S3.** Datasets used for the statistical analysis. The body mass index (BMI) and age values for each dataset are reported as an average value with minimum and maximum values in parentheses.

## References

1. Ley RE, Turnbaugh PJ, Klein S, Gordon JI. 2006. Microbial ecology: Human gut microbes associated with obesity. Nature 444:1022–1023.

2. Turnbaugh PJ, Ley RE, Mahowald MA, Magrini V, Mardis ER, Gordon JI. 2006. An obesity-associated gut microbiome with increased capacity for energy harvest. Nature 444:1027– 1031.

3. Waters JL, Ley RE. 2019. The human gut bacteria *Christensenellaceae* are widespread, heritable, and associated with health. BMC Biol 17:83.

4. Goodrich JK, Waters JL, Poole AC, Sutter JL, Koren O, Blekhman R, Beaumont M, Van Treuren W, Knight R, Bell JT, Spector TD, Clark AG, Ley RE. 2014. Human genetics shape the gut microbiome. Cell 159: 789–799.

5. Fu J, Bonder MJ, Cenit MC, Tigchelaar EF, Maatman A, Dekens JAM, Brandsma E, Marczynska J, Imhann F, Weersma RK, Franke L, Poon TW, Xavier RJ, Gevers D, Hofker MH, Wijmenga C, Zhernakova A. 2015. The gut microbiome contributes to a substantial proportion of the variation in blood lipids. Circ Res 117:817–824.

6. Goodrich JK, Davenport ER, Beaumont M, Jackson MA, Knight R, Ober C, Spector TD, Bell JT, Clark AG, Ley RE. 2016. Genetic determinants of the gut microbiome in UK Twins. Cell Host Microbe 19:731–743.

7. Kummen M, Holm K, Anmarkrud JA, Nygård S, Vesterhus M, Høivik ML, Trøseid M, Marschall H-U, Schrumpf E, Moum B, Røsjø H, Aukrust P, Karlsen TH, Hov JR. 2016. The gut microbial profile in patients with primary sclerosing cholangitis is distinct from patients with ulcerative colitis without biliary disease and healthy controls. Gut 66:611–619.

8. Lim MY, You HJ, Yoon HS, Kwon B, Lee JY, Lee S, Song Y-M, Lee K, Sung J, Ko G. 2016. The effect of heritability and host genetics on the gut microbiota and metabolic syndrome. Gut 66:1031–1038.

9. Oki K, Toyama M, Banno T, Chonan O, Benno Y, Watanabe K. 2016. Comprehensive analysis of the fecal microbiota of healthy Japanese adults reveals a new bacterial lineage associated with a phenotype characterized by a high frequency of bowel movements and a lean body type. BMC Microbiol 16:284.

10. Stanislawski MA, Dabelea D, Wagner BD, Sontag MK, Lozupone CA, Eggesbø M. 2017. Pre-pregnancy weight, gestational weight gain, and the gut microbiota of mothers and their infants. Microbiome 5:113.

11. Yun Y, Kim H-N, Kim SE, Heo SG, Chang Y, Ryu S, Shin H, Kim H-L. 2017. Comparative analysis of gut microbiota associated with body mass index in a large Korean cohort. BMC Microbiol 17:151.

12. Brooks AW, Priya S, Blekhman R, Bordenstein SR. 2018. Gut microbiota diversity across ethnicities in the United States. PLoS Biol 16:e2006842.

13. Jackson MA, Bonder MJ, Kuncheva Z, Zierer J, Fu J, Kurilshikov A, Wijmenga C, Zhernakova A, Bell JT, Spector TD, Steves CJ. 2018. Detection of stable community structures within gut microbiota co-occurrence networks from different human populations. PeerJ 6:e4303.

14. López-Contreras BE, Morán-Ramos S, Villarruel-Vázquez R, Macías-Kauffer L, Villamil-Ramírez H, León-Mimila P, Vega-Badillo J, Sánchez-Muñoz F, Llanos-Moreno LE, Canizalez-Román A, Del Río-Navarro B, Ibarra-González I, Vela-Amieva M, Villarreal-Molina T, Ochoa-Leyva A, Aguilar-Salinas CA, Canizales-Quinteros S. 2018. Composition of gut microbiota in obese and normal-weight Mexican school-age children and its association with metabolic traits. Pediatr Obes 13:381–388.

15. Peters BA, Shapiro JA, Church TR, Miller G, Trinh-Shevrin C, Yuen E, Friedlander C, Hayes RB, Ahn J. 2018. A taxonomic signature of obesity in a large study of American adults. Sci Rep 8:9749.

16. Morotomi M, Nagai F, Watanabe Y. 2012. Description of *Christensenella minuta* gen. nov., sp. nov., isolated from human faeces, which forms a distinct branch in the order Clostridiales, and proposal of Christensenellaceae fam. nov. Int J Syst Evol Microbiol 62:144–149.

17. Costea PI, Hildebrand F, Arumugam M, Bäckhed F, Blaser MJ, Bushman FD, Vos WM de, Ehrlich SD, Fraser CM, Hattori M, Huttenhower C, Jeffery IB, Knights D, Lewis JD, Ley RE, Ochman H, O’Toole PW, Quince C, Relman DA, Shanahan F, Sunagawa S, Wang J, Weinstock GM, Wu GD, Zeller G, Zhao L, Raes J, Knight R, Bork P. 2018. Enterotypes in the landscape of gut microbial community composition. Nature Microbiology 3:8–16.

18. Turpin W, Espin-Garcia O, Xu W, Silverberg MS, Kevans D, Smith MI, Guttman DS, Griffiths A, Panaccione R, Otley A, Xu L, Shestopaloff K, Moreno-Hagelsieb G, GEM Project Research Consortium, Paterson AD, Croitoru K. 2016. Association of host genome with intestinal microbial composition in a large healthy cohort. Nat Genet 48:1413–1417.

19. Bonder MJ, Kurilshikov A, Tigchelaar EF, Mujagic Z, Imhann F, Vila AV, Deelen P, Vatanen T, Schirmer M, Smeekens SP, Zhernakova DV, Jankipersadsing SA, Jaeger M, Oosting M, Cenit MC, Masclee AAM, Swertz MA, Li Y, Kumar V, Joosten L, Harmsen H, Weersma RK, Franke L, Hofker MH, Xavier RJ, Jonkers D, Netea MG, Wijmenga C, Fu J, Zhernakova A. 2016. The effect of host genetics on the gut microbiome. Nat Genet 48:1407–1412.

20. Hansen EE, Lozupone CA, Rey FE, Wu M, Guruge JL, Narra A, Goodfellow J, Zaneveld JR, McDonald DT, Goodrich JA, Heath AC, Knight R, Gordon JI. 2011. Pan-genome of the dominant human gut-associated archaeon, *Methanobrevibacter smithii*, studied in twins. Proc Natl Acad Sci USA 108 Suppl 1:4599–4606.

21. Upadhyaya B, McCormack L, Fardin-Kia AR, Juenemann R, Nichenametla S, Clapper J, Specker B, Dey M. 2016. Impact of dietary resistant starch type 4 on human gut microbiota and immunometabolic functions. Sci Rep 6:28797.

22. Klimenko NS, Tyakht AV, Popenko AS, Vasiliev AS, Altukhov IA, Ischenko DS, Shashkova TI, Efimova DA, Nikogosov DA, Osipenko DA, Musienko SV, Selezneva KS, Baranova A, Kurilshikov AM, Toshchakov SM, Korzhenkov AA, Samarov NI, Shevchenko MA, Tepliuk AV, Alexeev DG. 2018. Microbiome responses to an uncontrolled short-term diet intervention in the frame of the citizen science project. Nutrients 10:576.

23. Vanderhaeghen S, Lacroix C, Schwab C. 2015. Methanogen communities in stools of humans of different age and health status and co-occurrence with bacteria. FEMS Microbiol Lett 362:fnv092.

24. Moore WEC, Johnson JL, Holdeman LV. 1976. Emendation of *Bacteroidaceae* and *Butyrivibrio* and descriptions of *Desulfomonas* gen. nov. and ten new species in the genera *Desulfomonas*, *Butyrivibrio*, *Eubacterium*, *Clostridium*, and *Ruminococcus*. Int J Syst Evol Microbiol 26:238–252.

25. Traore SI, Khelaifia S, Armstrong N, Lagier JC, Raoult D. 2019. Isolation and culture of *Methanobrevibacter smithii* by co-culture with hydrogen-producing bacteria on agar plates. Clinical Microbiology and Infection 25:1561.e1–1561.e5.

26. Khelaifia S, Lagier J-C, Nkamga VD, Guilhot E, Drancourt M, Raoult D. 2016. Aerobic culture of methanogenic archaea without an external source of hydrogen. Eur J Clin Microbiol Infect Dis 35:985–991.

27. Nkamga VD, Lotte R, Roger P-M, Drancourt M, Ruimy R. 2016. *Methanobrevibacter smithii* and *Bacteroides thetaiotaomicron* cultivated from a chronic paravertebral muscle abscess. Clin Microbiol Infect 22:1008–1009.

28. Lau SKP, McNabb A, Woo GKS, Hoang L, Fung AMY, Chung LMW, Woo PCY, Yuen K-Y. 2007. *Catabacter hongkongensis* gen. nov., sp. nov., isolated from blood cultures of patients from Hong Kong and Canada. J Clin Microbiol 45:395–401.

29. Rosa BA, Hallsworth-Pepin K, Martin J, Wollam A, Mitreva M. 2017. Genome sequence of *Christensenella minuta* DSM 22607T. Genome Announc 5:e01451–16.

30. Hillmann B, Al-Ghalith GA, Shields-Cutler RR, Zhu Q, Gohl DM, Beckman KB, Knight R, Knights D. 2018. Evaluating the information content of shallow shotgun metagenomics. mSystems 3:e00069–18.

31. Gill SR, Pop M, Deboy RT, Eckburg PB, Turnbaugh PJ, Samuel BS, Gordon JI, Relman DA, Fraser-Liggett CM, Nelson KE. 2006. Metagenomic analysis of the human distal gut microbiome. Science 312:1355–1359.

32. Dridi B, Raoult D, Drancourt M. 2011. Archaea as emerging organisms in complex human microbiomes. Anaerobe 17:56–63.

33. Edwards T, McBride BC. 1975. Biosynthesis and degradation of methylmercury in human faeces. Nature 253:463–464.

34. Balch WE, Wolfe RS. 1976. New approach to the cultivation of methanogenic bacteria: 2-mercaptoethanesulfonic acid (HS-CoM)-dependent growth of *Methanobacterium ruminantium* in a pressurized atmosphere. Appl Environ Microbiol 32:781–791.

35. Stams AJM, Plugge CM. 2009. Electron transfer in syntrophic communities of anaerobic bacteria and archaea. Nat Rev Microbiol 7:568–577.

36. Shen L, Zhao Q, Wu X, Li X, Li Q, Wang Y. 2015. Interspecies electron transfer in syntrophic methanogenic consortia: From cultures to bioreactors. Renewable Sustainable Energy Rev 54:1358–1367.

37. Angenent LT, Karim K, Al-Dahhan MH, Wrenn BA, Domíguez-Espinosa R. 2004. Production of bioenergy and biochemicals from industrial and agricultural wastewater. Trends Biotechnol 22:477–485.

38. Macfarlane S, Macfarlane GT. 2003. Regulation of short-chain fatty acid production. Proc Nutr Soc 62:67–72.

39. Liu Y, Whitman WB. 2008. Metabolic, phylogenetic, and ecological diversity of the methanogenic *Archaea*. Ann N Y Acad Sci 1125:171–189.

40. den Besten G, van Eunen K, Groen AK, Venema K, Reijngoud D-J, Bakker BM. 2013. The role of short-chain fatty acids in the interplay between diet, gut microbiota, and host energy metabolism. J Lipid Res 54:2325–2340.

41. Louis P, Flint HJ. 2017. Formation of propionate and butyrate by the human colonic microbiota. Environ Microbiol 19:29–41.

42. Hoyles L, Swann J. 2019. Influence of the human gut microbiome on the metabolic phenotype, p. 535–560. In Lindon, JC, Nicholson, JK, Holmes, E (eds.), The Handbook of Metabolic Phenotyping. Elsevier.

43. Mbakwa CA, Penders J, Savelkoul PH, Thijs C, Dagnelie PC, Mommers M, Arts ICW. 2015. Gut colonization with *Methanobrevibacter smithii* is associated with childhood weight development. Obesity 23:2508–2516.

44. Morrison DJ, Preston T. 2016. Formation of short chain fatty acids by the gut microbiota and their impact on human metabolism. Gut Microbes 7:189–200.

45. Mack I, Cuntz U, Grämer C, Niedermaier S, Pohl C, Schwiertz A, Zimmermann K, Zipfel S, Enck P, Penders J. 2016. Weight gain in anorexia nervosa does not ameliorate the faecal microbiota, branched chain fatty acid profiles, and gastrointestinal complaints. Sci Rep 6:26752.

46. Armougom F, Henry M, Vialettes B, Raccah D, Raoult D. 2009. Monitoring bacterial community of human gut microbiota reveals an increase in *Lactobacillus* in obese patients and methanogens in anorexic patients. PLoS One 4:e7125.

47. Million M, Maraninchi M, Henry M, Armougom F, Richet H, Carrieri P, Valero R, Raccah D, Vialettes B, Raoult D. 2012. Obesity-associated gut microbiota is enriched in *Lactobacillus reuteri* and depleted in *Bifidobacterium animalis* and *Methanobrevibacter smithii*. Int J Obes 36:817–825.

48. Schwiertz A, Taras D, Schäfer K, Beijer S, Bos NA, Donus C, Hardt PD. 2010. Microbiota and SCFA in lean and overweight healthy subjects. Obesity 18:190–195.

49. Blaxter KL, Clapperton JL. 1965. Prediction of the amount of methane produced by ruminants. Br J Nutr 19:511–522.

50. Crutzen PJ, Aselmann I, Seiler W. 1986. Methane production by domestic animals, wild ruminants, other herbivorous fauna, and humans. Tellus B, 38B: 271–284.

51. Callaway TR, Edrington TS, Rychlik JL, Genovese KJ, Poole TL, Jung YS, Bischoff KM, Anderson RC, Nisbet DJ. 2003. Ionophores: their use as ruminant growth promotants and impact on food safety. Curr Issues Intest Microbiol 4:43–51.

52. Kruger Ben Shabat S, Sasson G, Doron-Faigenboim A, Durman T, Yaacoby S, Berg Miller ME, White BA, Shterzer N, Mizrahi I. 2016. Specific microbiome-dependent mechanisms underlie the energy harvest efficiency of ruminants. ISME J 10:2958–2972.

53. Poole AC, Goodrich JK, Youngblut ND, Luque GG, Ruaud A, Sutter JL, Waters JL, Shi Q, El-Hadidi M, Johnson LM, Bar HY, Huson DH, Booth JG, Ley RE. 2019. Human Salivary Amylase Gene Copy Number Impacts Oral and Gut Microbiomes. Cell Host Microbe 25:553–564.e7.

54. Pasolli E, Schiffer L, Manghi P, Renson A, Obenchain V, Truong DT, Beghini F, Malik F, Ramos M, Dowd JB, Huttenhower C, Morgan M, Segata N, Waldron L. 2017. Accessible, curated metagenomic data through ExperimentHub. Nat Methods 14:1023.

55. de la Cuesta-Zuluaga J, Ley RE, Youngblut ND. 2019. Struo: a pipeline for building custom databases for common metagenome profilers. Bioinformatics, btz899.

56. Wood DE, Salzberg SL. 2014. Kraken: ultrafast metagenomic sequence classification using exact alignments. Genome Biol 15:R46.

57. Lu J, Breitwieser FP, Thielen P, Salzberg SL. 2017. Bracken: estimating species abundance in metagenomics data. PeerJ Computer Science 3:e104.

58. Mende DR, Letunic I, Huerta-Cepas J, Li SS, Forslund K, Sunagawa S, Bork P. 2017. proGenomes: a resource for consistent functional and taxonomic annotations of prokaryotic genomes. Nucleic Acids Res 45:D529–D534.

59. Mangiafico SS. 2016. Summary and Analysis of Extension Program Evaluation in R, version 1.15.0.

60. R Core Team. 2017. R: A language and environment for statistical computing. R Foundation for Statistical Computing, Vienna, Austria.

61. Lambrecht J, Cichocki N, Hübschmann T, Koch C, Harms H, Müller S. 2017. Flow cytometric quantification, sorting and sequencing of methanogenic archaea based on F_420_ autofluorescence. Microb Cell Fact 16:180.

62. Schindelin J, Arganda-Carreras I, Frise E, Kaynig V, Longair M, Pietzsch T, Preibisch S, Rueden C, Saalfeld S, Schmid B, Tinevez J-Y, White DJ, Hartenstein V, Eliceiri K, Tomancak P, Cardona A. 2012. Fiji: an open-source platform for biological-image analysis. Nat Methods 9:676–682.

63. Karasov TL, Almario J, Friedemann C, Ding W, Giolai M, Heavens D, Kersten S, Lundberg DS, Neumann M, Regalado J, Neher RA, Kemen E, Weigel D. 2018. Arabidopsis thaliana and Pseudomonas Pathogens Exhibit Stable Associations over Evolutionary Timescales. Cell Host Microbe 24:168–179.e4.

64. Droop AP. 2016. fqtools: an efficient software suite for modern FASTQ file manipulation. Bioinformatics 32:1883–1884.

65. Jiang H, Lei R, Ding S-W, Zhu S. 2014. Skewer: a fast and accurate adapter trimmer for next-generation sequencing paired-end reads. BMC Bioinformatics 15:182.

66. Ewels P, Magnusson M, Lundin S, Käller M. 2016. MultiQC: summarize analysis results for multiple tools and samples in a single report. Bioinformatics 32:3047–3048.

67. Garcia-Betancur JC, Yepes A, Schneider J, Lopez D. 2012. Single-cell Analysis of Bacillus subtilis Biofilms Using Fluorescence Microscopy and Flow Cytometry. J Vis Exp 1–8.

68. Zhang H, DiBaise JK, Zuccolo A, Kudrna D, Braidotti M, Yu Y, Parameswaran P, Crowell MD, Wing R, Rittmann BE, Krajmalnik-Brown R. 2009. Human gut microbiota in obesity and after gastric bypass. Proc Natl Acad Sci U S A 106:2365–2370.

69. Turnbaugh PJ, Hamady M, Yatsunenko T, Cantarel BL, Duncan A, Ley RE, Sogin ML, Jones WJ, Roe BA, Affourtit JP, Egholm M, Henrissat B, Heath AC, Knight R, Gordon JI. 2009. A core gut microbiome in obese and lean twins. Nature 457: 480–484.

